# Loss of *Rnf43* accelerates *Kras*-mediated neoplasia and remodels the tumor immune microenvironment in pancreatic adenocarcinoma

**DOI:** 10.1101/2021.05.29.446305

**Authors:** Abdel Nasser Hosein, Gita Dangol, Takashi Okumura, Jason Roszik, Kimal Rajapakshe, Megan Siemann, Bidyut Ghosh, Maria Monberg, Paola A. Guerrero, Mohamed Zaid, Aatur Singhi, Cara L Haymaker, Hans Clevers, Sonja M. Woermann, Lotfi Abou-Elkacem, Anirban Maitra

**Affiliations:** Department of Translational Molecular Pathology, Ahmad Center for Pancreatic Cancer Research, The University of Texas MD Anderson Cancer Center, Houston, Texas, USA; Department of Internal Medicine, Division of Hematology & Oncology, The University of Texas Southwestern Medical Center, Dallas, Texas, USA; Advocate Aurora Health, Vince Lombardi Cancer Clinic - Sheboygan, Wisconsin, USA; Department of Melanoma Medical Oncology Research, Division of Cancer Medicine, The University of Texas MD Anderson Cancer Center, Houston, Texas, USA; Department of Radiation Oncology, The University of Texas MD Anderson Cancer Center, Houston, Texas, USA; Oncode Institute, Hubrecht Institute, Royal Netherlands Academy of Arts and Sciences, University Medical Center Utrecht and Princess Maxima Center, Utrecht, the Netherlands; Department of Pathology, University of Pittsburgh Medical Center, Pittsburgh, Pennsylvania, USA

## Abstract

*RNF43* is an E3 ubiquitin ligase that is recurrently mutated in pancreatic ductal adenocarcinoma (PDAC) and precursor cystic neoplasms of the pancreas. The impact of *RNF43* mutations on PDAC is poorly understood and autochthonous models have not been sufficiently characterized. In this study we describe a genetically engineered mouse model (GEMM) of PDAC with conditional expression of oncogenic *Kras* and deletion of the catalytic domain of *Rnf43* (KRC) in exocrine cells. We demonstrate that *Rnf43* loss results in an increased incidence of high-grade cystic lesions of the pancreas and PDAC. Importantly, KRC mice have a significantly decreased survival compared to mice containing only an oncogenic *Kras* mutation. By use of single cell RNA sequencing we demonstrated that KRC tumor progression is accompanied by a decrease in macrophages, as well as an increase in T and B lymphocytes with evidence of increased immune checkpoint molecule expression and affinity maturation, respectively. This was in stark contrast to the tumor immune microenvironment observed in the *Kras*/*Tp53* driven PDAC GEMM. Furthermore, expression of the chemokine, CXCL5, was found to be specifically decreased in KRC cancer cells by means of epigenetic regulation and emerged as a putative candidate for mediating the unique KRC immune landscape. This GEMM establishes *RNF43* as a *bona fide* tumor suppressor gene in PDAC and puts forth a rationale for an immunotherapy approach in this subset of PDAC cases.

## Introduction

Pancreatic ductal adenocarcinoma (PDAC) is the third leading cause of cancer-related deaths in the United states, and an estimated 48,220 people will succumb to the disease in 2021 [1]. The incidence of PDAC is on the rise, and PDAC is projected to become the second most common cause of cancer death in the USA by 2030 [2]. Furthermore, 80-85% of patients present with advanced disease and therefore are not eligible for curative intent surgery [3]. As a result, there is a critical need for precision based systemic treatment options are needed to address this growing public health problem. To that end, whole exome and whole genome sequencing efforts have revealed a compendium of recurrent mutations in human PDAC [4–6]. Specifically, 5-10% of resected PDAC cases display loss of function mutations in Ringer Finger Protein 43, *RNF43*. In addition, *RNF43* has been shown to be one of the most commonly mutated genes in pancreatic cystic neoplasms, including intraductal papillary mucinous neoplasms (IPMN) and mucinous cystic neoplasms, which are *bona fide* precursors to PDAC [7, 8]. *RNF43* is also frequently mutated in several other cancer types such as colon and endometrial adenocarcinomas [9].

*RNF43*, an E3 ubiquitin ligase enzyme, uses a ring finger domain to catalyze the final step of protein ubiquitination; notably, it has been demonstrated to ubiquitinate Frizzled receptors at the plasma membrane and, in doing so, acts as a negative regulator of Wnt signaling [10]. Recently, preclinical models have demonstrated targeted vulnerability of *RNF43* mutated human PDAC cell lines to blockade of specific Frizzled receptors [11] and small molecule porcupine inhibition, an enzyme critical to the production of functional Wnt ligands [12]. Despite these promising preclinical findings, however, a phase 1 clinical trial of the porcupine inhibitor, WNT974, in *RNF43* mutated solid cancer patients did not display any signal of activity [13]. As such, improved preclinical models of *RNF43* mutated PDAC are needed, both to better elucidate the biology of how this mutation impacts *Kras*-induced pancreatic neoplasia, and further, for the development of precision-based therapeutics for this subset of patients.

In this report we describe a genetically engineered mouse model (GEMM) of PDAC driven by oncogenic *Kras* and loss of the ring finger domain of *Rnf43*. We demonstrate that *Rnf43* is indeed a *bona fide* tumor suppressor gene in PDAC, as its conditional deletion cooperates with oncogenic *Kras* to increase the incidence of high-grade cystic lesions and subsequent PDAC. Remarkably, we also show that loss of *Rnf43* leads to a distinct tumor immune microenvironment (TIME), characterized by the preponderance of adaptive immune cells at the expense of immunosuppressive myeloid cells, when compared to the prototypal “*Kras;Tp53*” (“KPC”) mutant PDAC GEMMs with intact *Rnf43* function. This characteristic TIME in the setting of *Rnf43* mutations open up the possibility of utilizing immune checkpoint inhibitors as a potential therapeutic strategy in this subset of patients, in a disease which is essentially recalcitrant to immunotherapy. Our study also reiterates the importance of autochthonous organisms in identifying therapeutic dependencies that might otherwise not be uncovered in an *ex vivo* or hetero-transplant model context.

## Results

### Deficiency of *Rnf43* accelerates the development of *Kras-*driven precursor lesions

To investigate the role of *Rnf43* in pancreatic cancer development, we crossed mice bearing *Ptf1a-Cre* and LSL-*Kras*^G12D^ alleles with those bearing *Rnf43^flox^* alleles to generate compound progeny with pancreas specific mutant *Kras* expression, and either a heterozygous loss of *Rnf43* in the pancreas (KRC^het^) or with bi-allelic loss of *Rnf43* in the pancreas (KRC) (**Figure 1A**). *Cre*-mediated recombination of *LSL-Kras^G12D^* allele in the mouse pancreas activates oncogenic *Kras*, which is sufficient to develop pancreatic precursor lesions, and in a subset, results in adenocarcinomas [14]. At 9 weeks of age, both KRC^het^ and KRC mice developed enlarged pancreata compared to *Ptf1a-Cre;Kras^G12D^* (KC) mice (**Figure 1B**). Quantitative analysis of pancreatic tissues in KRC^het^ and KRC mice at nine weeks old showed significantly greater areas of acinar parenchyma loss in comparison to age matched KC mice (**Figure 1C**). This was accompanied by an increase in ductal precursor lesions, delineated by expression of the ductal marker SOX9, which also displayed an increased proliferative index with greater Ki67 staining in both the KRC^het^ and KRC pancreata compared to KC mice (**Figures 1D and 1E**).

**Figure 1:**
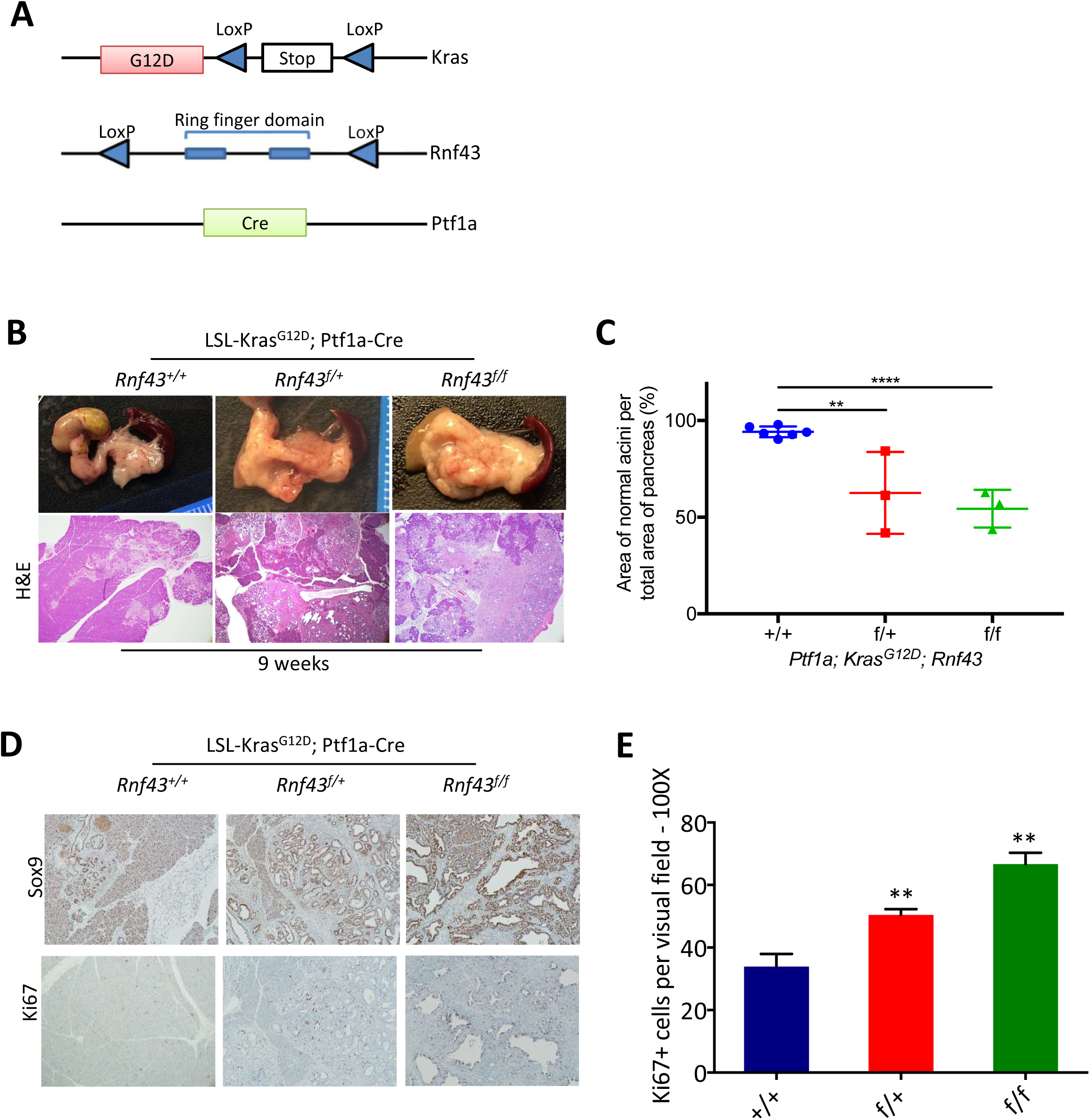
**Deficiency of *Rnf43* accelerated the development of *Kras-driven*** pancreatic ductal precursor lesions. (A) Schematic of strategy used to generate mouse models with pancreas-specific conditional knockout of *Rnf43* and Kras^G12D^ mutation; (B) Gross anatomy (top) and H&E stained sections (bottom) of mouse pancreata from respective genotypes showing precursor lesions of pancreas (20x); (C) Graph representing the percentage of normal acinar area over total area of pancreas from each respective genotype. (D) Immunohistochemistry showing expression of SOX9 (10X) and Ki67(10X) in pre-malignant lesions in pancreata from the indicated genotypes. (E) Quantitative analysis of Ki67 positive cells in precursor lesions from respective genotypes. 10 visual fields per mice and three mice per genotype were analyzed. Unpaired t-test was performed. Error bars, s.e.m. **p<0.01

Regarding the role of *Rnf43* in pancreatic development and tumorigenesis, we noted that loss of *Rnf43* is dispensable for normal pancreas development as pancreas specific loss of *Rnf43* in the absence of an activating *Kras* mutation (RC^het^ or RC mice) resulted in normal gross and histological morphology of pancreatic tissue (**Suppl Figure 1A, B**). We also noted that immunohistochemical (IHC) analyses of amylase showed mature acinar cell architecture, while IHC for glucagon and insulin also demonstrated no abnormalities in islets cells. (**Supp Figure 1B**). Pancreatic weight:body weight ratios were equivalent across KC, RC^het^ and RC mice (**Supp Figure 1C**) at 9 weeks of age. Importantly, during the course of observation, up to the age of 62 weeks of age, none of the RC^het^ or RC mice developed pancreatic neoplasia; both RC^het^ and RC mice demonstrated normal pancreatic tissue architecture with no evidence of gross anatomical, histologic, or immunohistochemical abnormalities at all time points examined (RC^het^, n=17, 9-62 weeks; RC, n=24, 9-61 weeks) (**Supp Figure 1D**). These data indicate that loss of *Rnf43* is dispensable for pancreas development and requires the presence of oncogenic *Kras* in order to initiate pancreatic neoplasia.

### Loss of *Rnf43* cooperates with oncogenic *Kras* in development of murine cystic precursor lesions

*RNF43* is recurrently mutated in mucinous cystic precursor lesions of the pancreas in humans, and most also harbor concurrent oncogenic *KRAS* mutation [8]. Histological examination of pancreatic tissue of mice below the age of 52 weeks (including timed necropsy and survival cohorts), demonstrated significantly higher prevalence of cystic precursor lesions resembling IPMNs in KRC mice (41%) compared to KRC^het^ mice (15%) and KC mice (5%) (**Figure 2A-B**). KRC mice developed high grade cystic precursor lesions as early as nine weeks, a finding not observed in either KC or KRC^het^ (**Supp Figure 1A**), while the both KRC^het^ and KRC demonstrated a significant increase in low grade cystic lesions compared to KC mice at nine weeks old (**Supp Figure 2B**). Pancreatic cystic lesions were appreciated on gross morphology and recapitulated cystic histology seen in human IPMN (**Supp Figure 2C**). By five months of age, both the KRC^het^ and KRC demonstrated an increase in the incidence of high-grade lesions relative to pancreata from the KC cohort (**Supp Figure 2D**). These data demonstrate the ability of *Rnf43* loss in the KRC GEMM to cooperate with *Kras* in producing cystic lesions of the pancreas akin to *RNF43* loss of function mutations in human patient cohorts.

**Figure 2:**
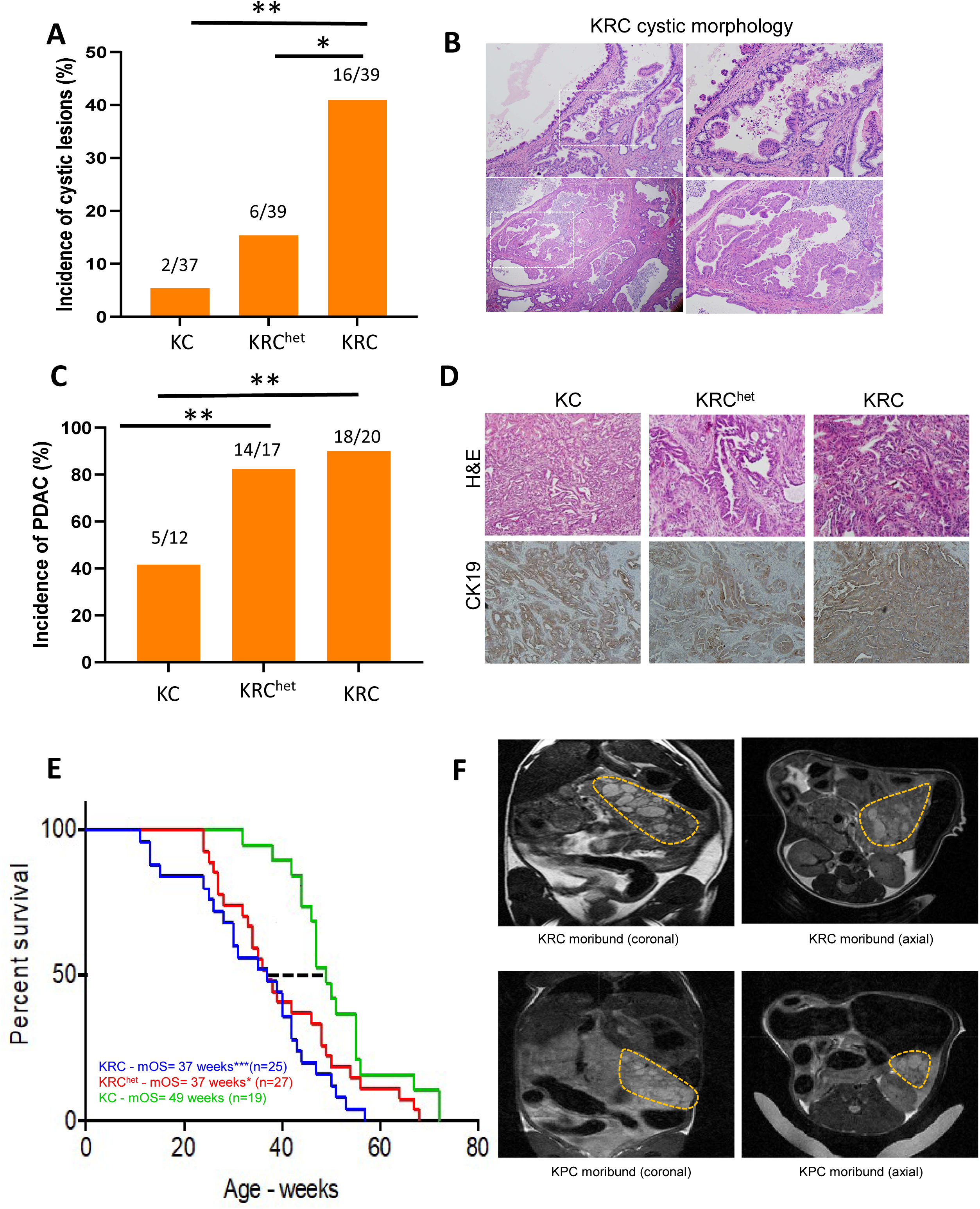
**Loss of *Rnf43* cooperates with oncogenic *Kras* in the development of cystic lesions and invasive ductal adenocarcinoma of the pancreas with shortened survival.** (A) Graph displaying the incidence of cystic lesions in mice below the age of 52 weeks, including mice from both survival and timed-necropsy cohorts. Fisher’s exact test was performed to calculate p-value. (B) Representative images of cystic histology showing mucinous papillary projections of ductal epithelium. Left panel 4X; outlined section enlarged on right panel, 20X. (C) Graph showing cancer incidences of mice within the age of 52 weeks. (D) Representative images of H&E and CK19 staining of PDAC from indicated genotypes (20X magnification). (E) Overall survival analysis of mice with indicated genotypes. Log-rank test was used to evaluate the significance of survival differences between indicated genotypes. (*p <0.05; ***p<0.001). (F) T2 weighted dynamic contrast enhanced MR of a moribund KRC mouse (top) and KPC mouse (bottom). Coronal (left) and axial (right) images from the same imaging study are displayed.

### *Pancreas specific loss of Rnf43* cooperates with oncogenic *Kras* to mediate the development of pancreatic adenocarcinoma and decrease survival

To determine the phenotypic impact of *Rnf43* loss in the context of oncogenic *Kras*, we analyzed the incidence of PDAC in mice of the 3 different genotypes. Overall, we observed a higher incidence of invasive cancer in timed necropsies until 52 weeks of age in KRC^het^ and KRC pancreata compared to KC mice (**Figure 2C-D**). From our histological analyses of the resulting murine PDAC, we could not delineate with certainty if the PDAC arose from a pre-existing cystic precursor, although we did observe invasive adenocarcinomas even in pancreata with no obvious residual cystic lesions. Notably, *RNF43* mutations are identified in “conventional” human PDAC, without associated IPMNs or MCNs [6], which is consistent with our findings. We also evaluated our cohorts to determine the status of metastatic lesions across different genotypes. We also observed that a minority of KRC mice had metastatic disease at the time of necropsy with ∼21% having lung metastases and ∼27% having liver metastases. Of note, metastatic disease could be observed on T2 weighted MRI (**Supp Figure 3**). Finally, we performed survival analyses in cohorts of KC, KRC^het^ and KRC mice in which mice were euthanized when they became moribund. Kaplan-Meier survival analysis revealed a significant reduction in survival accompanied by loss of one or both *Rnf43* alleles, with a median survival of 37 weeks for both the homozygous KRC mice (p-value = 0.0003) and heterozygous KRC^het^ mice (p-value = 0.0231) compared to 49 weeks for *Rnf43* wild type KC mice (**Figure 2E**). Lastly, as part of our initial characterization of this novel GEMM we performed contrasted magnetic resonance imaging (MRI) of the moribund KRC GEMM and moribund KPC (*Ptf1a-Cre;Kras^G12D^*Tp53^R172H/+^) GEMM, whose genotype represents the majority of PDAC patients [6, 15]. On T2 weighted MRI intra-tumoral septations were readily appreciated in the cystic-solid KRC GEMM whereas the solid KPC GEMM featured a hazier and less well delineated tumor (**Figure 2F**)

### Single cell RNA-seq profiling of the KRC GEMM reveals progression toward a myeloid-low, lymphocyte-rich microenvironment

Single-cell RNA-sequencing (scRNAseq) was performed on freshly harvested KRC GEMM pancreata, in order to deeply interrogate alterations in the tumor microenvironment from early stages of disease to moribund status. The single-cell suspensions of KRC pancreata were analyzed by the 10X Genomics platform at an early stage (2 months, n=2), an intermediate stage (4 months, n=2), and moribund status (6-10 months, n=3). “Early” KRC (**Figure 3A, left**), “intermediate” KRC (**Figure 3A, center**) and “moribund” KRC (**Figure 3A, right**) contained 2774, 5488 and 3249 cells, respectively. For each UMAP, graph-based population delineation was performed and populations were assigned an identity based on lineage markers we and others have previously employed in assigning cell populations in PDAC GEMMs (**Figure 3B**) [16, 17]. Percentages of each cell type in each of the three time points were then summarized (**Figure 3C**).

**Figure 3:**
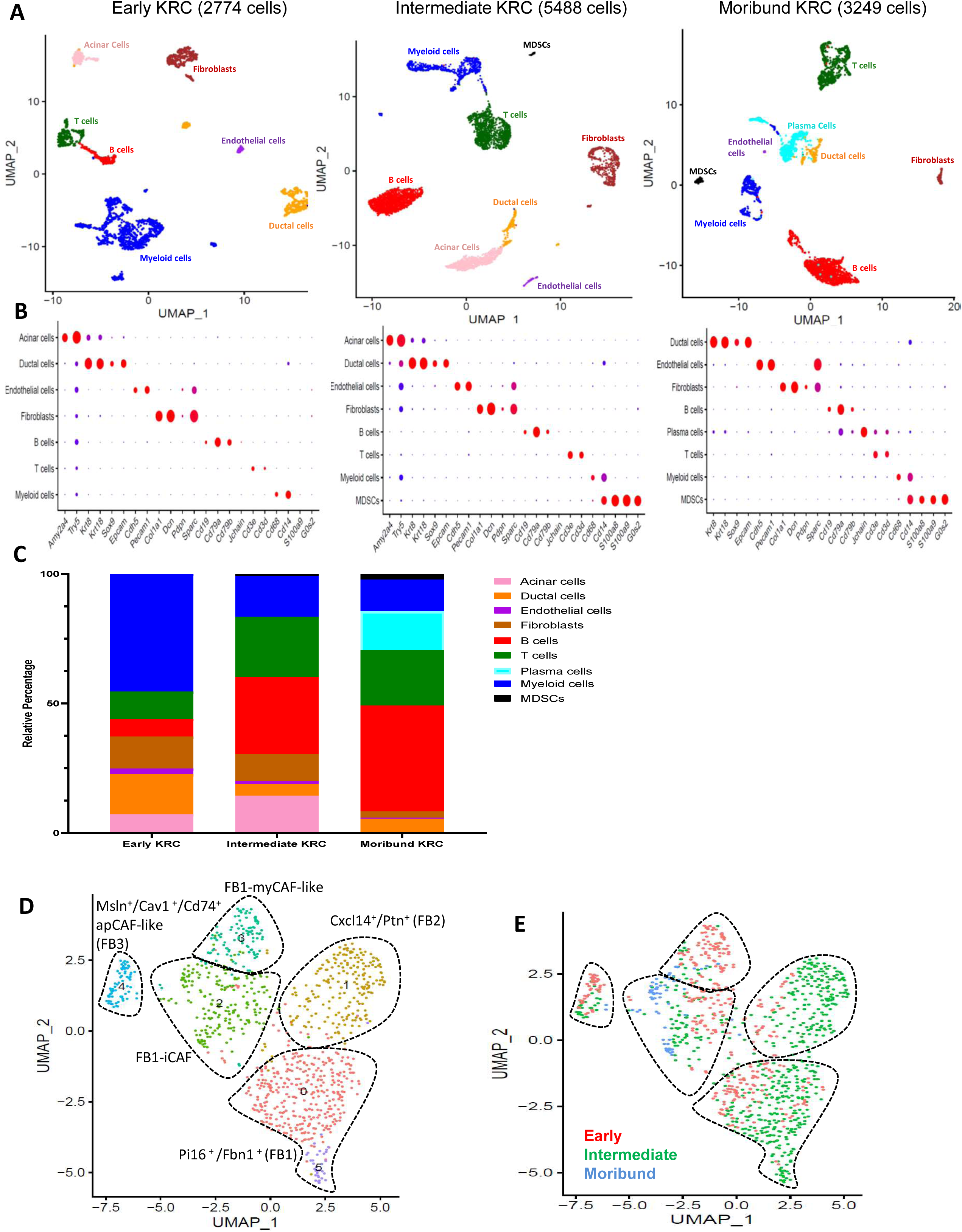
**scRNAseq analysis of KRC GEMM progression.** (A) UMAP plots of all cells in the early (left; 2774 cells), intermediate (center; 5488 cells) and moribund (right; 3249 cells) KRC GEMM stages. Cell lineage is labeled and denoted by color on each UMAP plot. (B) Dot plots corresponding to each of the UMAPs; KRC early (left), KRC intermediate (center), KRC moribund (right). (C) Corresponding representative H&E stained pancreata from each of the KRC stages; KRC early (left), KRC intermediate (center), KRC moribund (right). (D) Summary bar graph denoting the percentage of each cell type observed in the UMAPs for each of the three stages analyzed. Color legend appears in the graphic. D) UMAP displaying all fibroblasts observed across the three KRC stages (early, intermediate, moribund) with a graph-based population delineation method applied to illustrate fibroblast subpopulations. (E) Fibroblast UMAP with each cell color coded according to KRC stage (early = red, intermediate = green, moribund = blue).

We observed a diminishing percentage of fibroblasts during KRC progression, with fibroblasts comprising 12.3% of cells in early KRC, 10.3% of cells in intermediate KRC and 2.4% of cells in moribund KRC. This is a similar pattern as we previously observed in an oncogenic *Kras*/*Ink4a* loss driven PDAC GEMM (KIC), where the early time point featured greater than 3 times as many fibroblasts as the moribund time point by cellular percentage [16]. We then interrogated the KRC GEMM fibroblast populations in greater detail by collating and displaying the fibroblasts from each of the early, intermediate and moribund experiments on a single fibroblast UMAP (**Figure 3D**). We delineated the fibroblast populations based on scRNASeq expression clusters and noted the presence of a *Pi16^+^/Fbn1^+^* expressing population (Fibroblast 1; FB1), a *Cxcl14^+^/Ptn^+^* expressing fibroblast population (Fibroblast 2; FB2) and a *Cav1^+^/Msln^+^/Cd74^+^* expressing fibroblast population (Fibroblast 3; FB3) (**Supp Fig 4A**), reproducing the same three fibroblasts subtypes we had previously described [16]. In contrast to the presence of all three fibroblast subtypes in early and intermediate KRC, we noted that nearly all the KRC moribund fibroblasts belonged to only two distinct subpopulations; FB1-iCAF and FB1-myCAF-like [18], both of which had retained *Pi16^+^/Fbn1^+^* expression with no appreciable *Cxcl14^+^/Ptn^+^* expression (**Figure 3E**). Indeed, we had previously reported that all three fibroblast populations are present in normal mouse pancreas and early PDAC GEMMs, but FB2 fades by the moribund stage in two different PDAC GEMMs [16]. Moreover, in addition to *Pi16^+^/Fbn1^+^* expression, the FB1-iCAF population expressed several inflammatory cytokines (examples: *Il6* and *Ccl7*), similar to the inflammatory cancer associated fibroblast (iCAF) population, which has recently been described (**Supp Fig 4B**) [18]. The FB1-myCAF-like population contained *Acta2* and *Tagln* expressing cells, comparable to the myCAF population described by the same investigators. The FB3 population (comparable to apCAF) was present in early KRC and intermediate KRC, consistent with our previous reports in the KIC GEMM [16] however, these cells were greatly diminished by the moribund stage, in conjunction with an overall reduction in the cumulate CAF population.

Myeloid cells were noted to comprise the plurality of cells in the early KRC data set (45.5%) compared to intermediate KRC (15.7%) and moribund KRC experiments (12.3%) where the percentage of myeloid cells was approximately one third of that observed in early KRC (**Figure 3A-C**). Conversely, the early KRC GEMM contained only 10.6% cells as T cells whereas this percentage was greater than doubled in the intermediate (23.2%) and moribund (21.4%) KRC GEMMs. Importantly, we also noted a significant increase in T-cells present in *RNF43* human PDAC samples in comparison to *RNF43* wild type human PDAC samples (**Supp Figure 5A**). Most prominent was the expansion of the B cell lineage during KRC progression, which increased from KRC early (6.8%), to KRC intermediate (29.8%) and finally KRC moribund (40.9%). Consistent with this B cell expansion during KRC progression, we observed the emergence of a readily discernible plasma cell population by the KRC moribund stage which comprised ∼15% of cells; this latter phenomenon appears to be a unique feature of the KRC GEMM, as previous scRNAseq profiles on other clinically relevant PDAC GEMMs have not demonstrated a substantial plasma cell population [16, 17]. Together, B cell lineage (B lymphocytes + plasma cells) comprised greater than half of all cells by cellular percentage in the moribund KRC GEMM scRNAseq dataset. The intermediate KRC time point also noted the emergence of a population of myeloid-derived suppressor cells (MDSCs; ∼1%), which overexpressed *S100A8*, *S100A9* and *G0s2*, and expanded to 2.2% by the KRC moribund stage.

The immune composition observed in the moribund KRC GEMM featured a low myeloid:lymphocyte ratio and represents the inverse of the scenario we previously described in moribund KPC GEMM which featured a high myeloid:lymphocyte ratio (**Supp Table 1**). Interestingly, whereas we report a median survival of 37 weeks for the KRC GEMM, the previously published median survival of the *Kras/Tp53* driven PDAC GEMMs is of 12-20 weeks [19]. This survival discrepancy may, at least in part, be due to the decreased myeloid cells and increased lymphocytes seen in the KRC GEMM.

### KRC GEMM progression is accompanied by a decrease in inflammatory macrophages and preservation of monocytes and dendritic cells in the TIME

We next performed a focused myeloid lineage analysis, compiling all myeloid cells from the three stages of KRC GEMM progression and displaying them on a single UMAP plot (**Figure 4A, Supp Fig 6A**). These analyses revealed the presence of three distinct populations of dendritic cells; conventional dendritic cell 1 (cDC1) cells, which specifically expressed *Itgae, Irf8, Flt3, Clec9a, Xcr1*; second, cDC2 cells, which expressed *Cd209a*, and third a primed population of cDC1 cells (cDC1’), which like cDC1 expressed *Irf8* and *Flt3* but also expressed *Ccr7* and *Ccl5*. The latter are markers of cDC1 cells that have been primed by antigen in regional lymph nodes and has been described in the KPC GEMM [20]. Monocyte and macrophage cell populations were subsequently defined in the KRC GEMM. Importantly, in the KPC GEMM, it has been previously reported that embryonal derived, pancreatic tissue resident macrophages, expand during PDAC progression, and have pro-tumorigenic properties, while a monocytic population is recruited from the bone marrow during tumorigenesis and has a role in antigen presentation [21]. In our KRC scRNAseq data we assigned myeloid population identities based on previously described cell type markers [16, 17, 22, 23] and observed the presence of a large macrophage population that was comprised of 3 macrophage subpopulations. These macrophages were rich in inflammatory molecules such as *Ccl6, Il6* and the complement molecules *C1qa* and *C1qb*. Specifically, subpopulation 1 overexpressed the tissue resident macrophage marker *Adgre1* (F4/80), in addition to other inflammatory markers such as *Apoe*, *Pf4* and *Ccl3*. Of note, we observed a distinct *Cd14^+^/Thbs1^+^* monocyte population.

**Figure 4:**
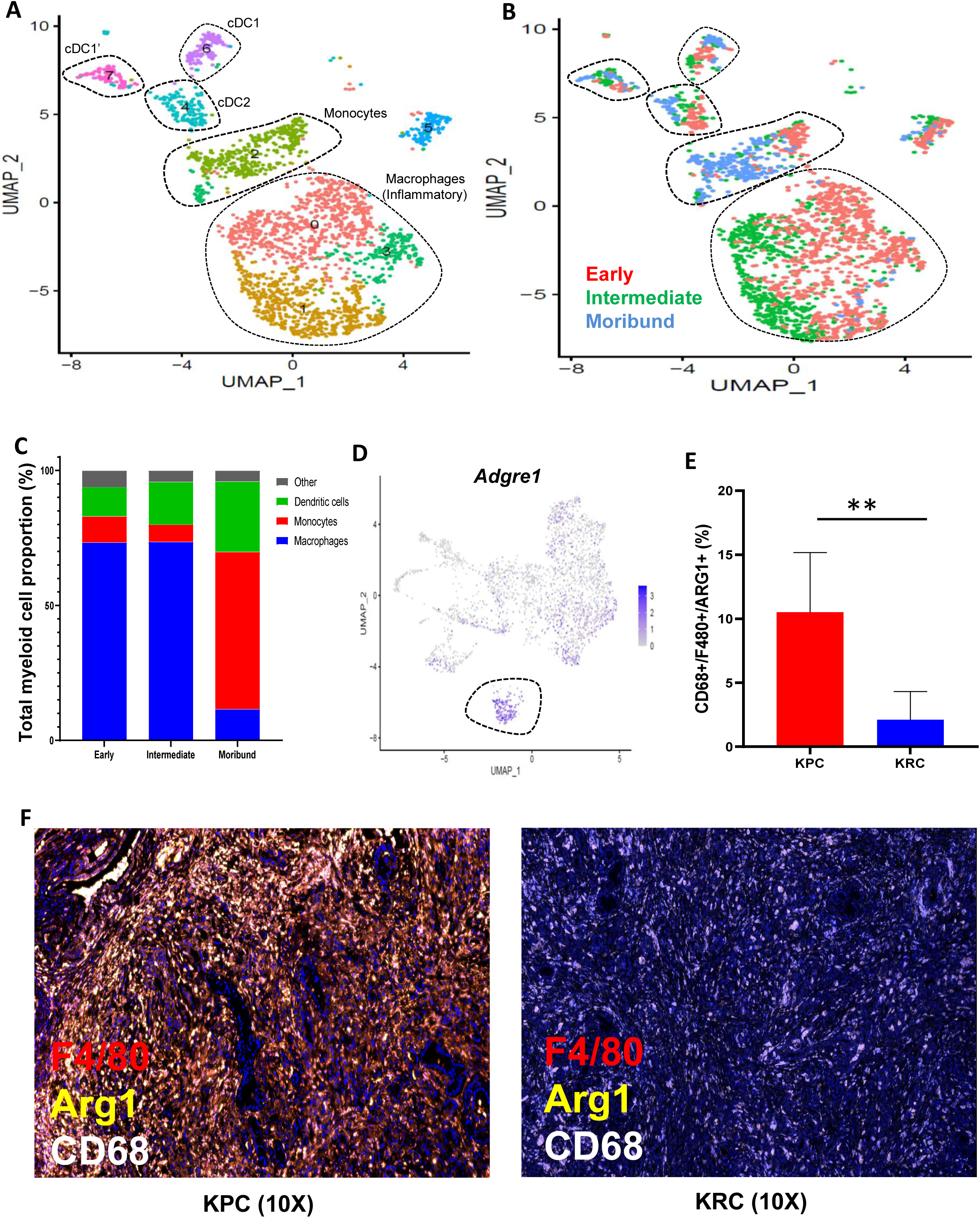
**Myeloid focused scRNAseq analysis of the KRC GEMM.** (A) UMAP displaying all cells of myeloid lineage observed across the three KRC stages (early, intermediate, moribund) with a graph-based population delineation method applied to illustrate myeloid subpopulations. (B) Myeloid UMAP with each cell color coded according to KRC stage (early = red, intermediate = green, moribund = blue). (C) Summary bar graph denoting the percentage of each myeloid cell type observed in each of the three KRC stages analyzed. (D) Single gene UMAP of all myeloid cells in KPC and KRC GEMMS with *Adgre1* expression denoted. The tissue resident macrophage population is outlined. (E) Quantification of CD68, ARG1 and F4/80 immunofluorescence in KRC (n=4) and KPC (n=3) tumor tissue sections. The percentage of cells positive for all three markers were averaged and displayed (two-sided t-test, *p=0.02). (F) Three color immunofluorescence analysis of macrophage markers, CD68 (white), ARG1 (gold), F4/80 (red) and DAPI (blue), in KPC (left) and KRC (right) tumors. Both images are displayed at 10X magnification

We then applied KRC stage to the myeloid lineage UMAP to determine if the prevalence of different myeloid cell types change during KRC GEMM progression (**Figure 4B**). Remarkably, we noted a substantial diminishment of pro-inflammatory/tissue resident macrophages in the KRC moribund stage, where these cells comprised only 11.5% of all myeloid cells, in comparison to the KRC early and intermediate stages where they comprised as much as 73.3% and 73.5% of all myeloid cells, respectively (**Figure 4C**). Monocytic cells and all 3 populations of dendritic cells were present across the 3 KRC GEMM stages making the marked decrease in tissue resident/inflammatory macrophages in the KRC moribund stage largely responsible for the overall marked reduction of myeloid cells in the moribund stage (**Figure 3D**). We also performed a comparative analysis of scRNAseq data of myeloid cell distribution from KRC moribund mice compared with myeloid cell distribution from moribund KPC mice we had previously reported [16]. By collating the myeloid cells from these two GEMMs (**Supp Fig 6B**) we observed a population of *Adgre1+* tissue resident macrophages in the KPC GEMM which is essentially devoid in the moribund KRC tumors (**Figure 4D**). We also noted a decrease in expression of inflammatory cytokines in KRC macrophages in comparison to KPC macrophages (**Supp Fig 5C**). We confirmed the marked decrease in F4/80^+^ (*Adgre1*) macrophages in KRC tumors relative to KPC tumors by multi-color immunofluorescence (**Figure 4E, F, G; Supp Fig 7**). The noticeable divergence in myeloid cell composition in moribund KRC *versus* KPC mice underscores the influence of *Kras* cooperating mutations in framing the respective TIME, and the possibility of leveraging this unique milieu for therapeutic benefit.

### Focused B-cell lineage analysis reveals expansion and clonal maturation of B cells during KRC progression

We pooled all B-cell lineage cells (B-cells and plasma cells) from the KRC early, intermediate and moribund experiments and displayed them on a single UMAP (**Figure 5A**). This analysis resulted in the segregation of cells into 3 distinct groups; B cells, early plasma cells (1 subgroup) and mature plasma cells (2 subgroups). As expected, based on previous analyses (**Figure 3)**, when the KRC stage identity was applied we noted that nearly all the plasma cells were derived from the KRC moribund stage (**Figure 5B**). Moreover, the early KRC B-cells were positioned toward the bottom of the UMAP with intermediate and ultimately moribund KRC B-cells showing a pseudo-trajectory toward regions of the plot in closer proximity to the plasma cells. We also performed clustering analyses using expression values of all significant immunoglobulin transcripts in the 3 subpopulations of plasma cells (**Figure 5C**). Theses analyses illustrated the specific over-expression of 2-3 immunoglobulin genes in each of the mature plasma cells subpopulations. Both mature plasma cell populations specifically over-expressed at least one variable heavy chain immunoglobulin gene and one variable light chain immunoglobulin gene. This is in contrast to the early plasma cell population which over-expressed *Jchain* and *Ighm*, which are markers of an early humoral response [24]. These data were also illustrated with single gene UMAPs of a gene specific for each of the late plasma cell and the early plasma cell populations (**Figure 5D**). When all B-cell lineage derived cells from the KRC moribund and KPC moribund GEMMs were collated on a single UMAP, we observed only a few cells in the main B lymphocyte population, in addition to the early plasma cell population, that were derived from the KPC mice, while the late plasma cell populations were almost entirely derived from the KRC tumors (**Supp Figure 8**). Taken together, these data indicate that, in sharp contrast to the KPC mice, disease progression in the KRC GEMM is uniquely accompanied by the expansion and maturation of an adaptive B-cell response, culminating in clonal plasma cells which have likely undergone affinity maturation possibly akin to what has been described in other solid tumors [25].

**Figure 5:**
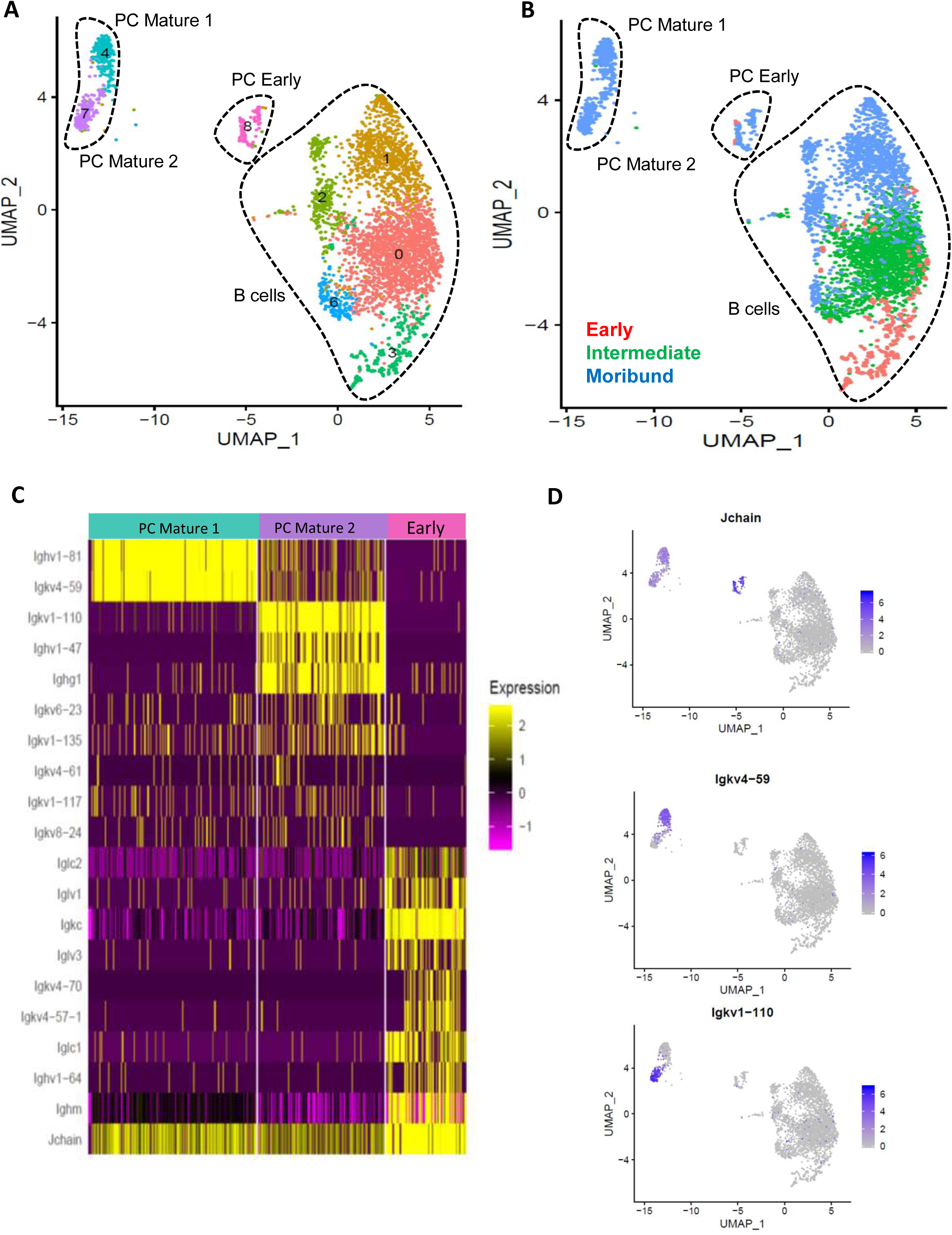
**B cell lineage focused scRNAseq analysis of the KRC GEMM.** (A) UMAP displaying all cells of B cell lineage observed across the three KRC stages (early, intermediate, moribund) with a graph-based population delineation method applied to illustrate B lineage subpopulations. (B) B lineage UMAP with each cell color coded according to KRC stage (early = red, intermediate = green, moribund = blue). (C) Immunoglobulin (Ig) focused heatmap displaying Ig gene expression across the two mature and one early plasma cell subpopulations. (D) Single gene UMAP displaying specific markers for the early plasma cells (top; *Jchain*), mature plasma cell 1 (middle; *Igkv4-59*), and mature plasma cell 2 (bottom; *Igkv1-110*).

### Focused T cell analysis of the KRC GEMM reveals regulatory T-cell immune checkpoint molecule upregulation as a potential mechanism of immune evasion during KRC progression

To further interrogate the temporal composition of the KRC TIME, all T-cells from the early, intermediate and moribund KRC scRNAseq datasets were projected on a single, T cell focused, UMAP (**Figure 6A**). Graph based population delineation identified commonly described T-cell subsets: naïve T-cells (*Ccr7, Lef1, Dusp10*), activated T-cells (*Cd8*, *Gzma, Gzmb, Ccl5*), regulatory T cells (Treg) and group 2 innate lymphoid cells (ILC2; *Gata3, Rora, Areg, Il5*) which has recently been shown to have a critical role in mediating PDAC sensitivity to immune checkpoint blockade [26]. We also observed the presence of dysfunctional T-cells that express *Id2, Cd8, Cd8b1*, and have recently been described in other immune checkpoint sensitive solid cancers (melanoma, non-small cell lung cancers) to be a major proliferating immune cell component [27]. When the KRC stage identities were overlayed with the T-cell UMAP (**Figure 6B**) we noted that the dysfunctional T cell compartment was composed entirely of cells derived from the KRC intermediate stage, indicating a proliferative population as the TIME shifts to a lymphocyte rich state. None of the naïve T-cells, T_regs_ or activated T-cells displayed dominance or depletion of any particular stage during stages of KRC GEMM progression. To investigate potential mechanisms of immune escape in the KRC GEMM we performed a focused analysis on all immune checkpoint or costimulatory genes expressed in the T_reg_ cell population (**Figure 6C**): *Ctla4, Pdcd1, Icos, Tnfsf4* (OX40)*, Tnfsf8* (CD153/CD30L) *and Cd83.* These analyses revealed an increase in expression intensity and percent of the T_reg_ population expressing each of these immune checkpoint genes as the KRC GEMM progressed from early to intermediate and moribund stages. A notable exception to this was *Icos* which demonstrated a decrease in expression intensity as the GEMM progressed, which is consistent with the function of ICOS as a co-stimulatory receptor in T-cell activation [28]. In light of this unique T cell contexture in the moribund KRC mice, we performed a proof-of-concept assay in which KRC mice were treated twice per week with either an anti-CTLA4 blocking antibody (n=6) or an IgG isotype control antibody (n=6), given by intraperitoneal injection for a fixed 24-day duration. CTLA4 was chosen as a target given its high expression in KRC moribund mice and its current clinical use as therapeutic immune checkpoint target in other solid cancers [29–31]. KRC mice were enrolled onto treatment at 6 months of age, randomized to one of the two arms and a baseline MRI was performed (**Figure 6D**). Imaging was repeated halfway through the trial and at the end of the treatment course. The primary endpoint of this assay was progression on imaging or mouse death while on treatment. During this trial, 4/6 mice in the IgG isotype treated group died while 2/6 mice in this group showed progression on imaging; all mice in this arm met the primary endpoint (**Figure 6E, F**). In contrast, 4/6 mice in the anti-CTLA4 antibody treated group experienced radiographic stability and 2/6 mice experienced a radiographic response; none of the mice in this arm met the primary endpoint.

**Figure 6:**
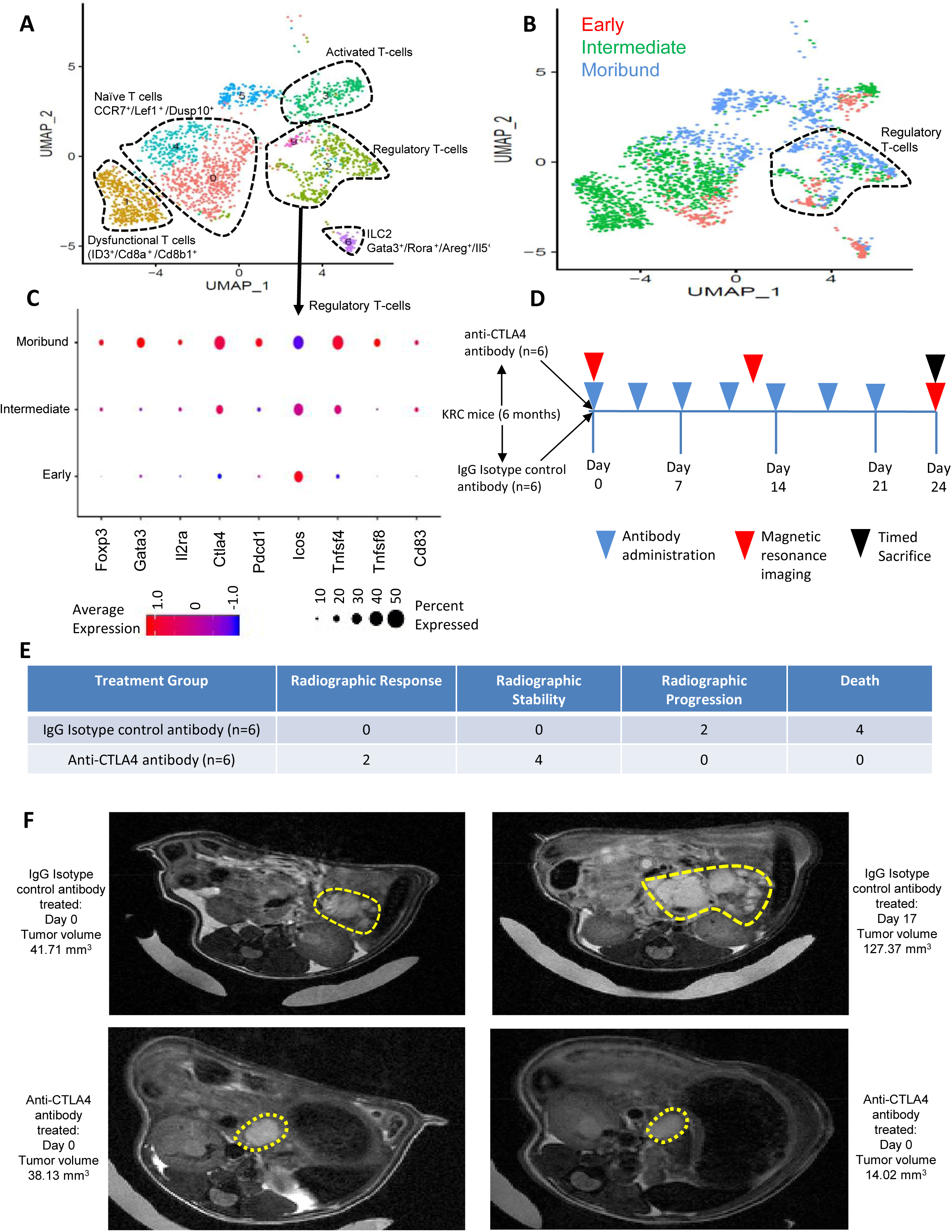
**T cell lineage focused scRNAseq analysis of the KRC GEMM.** (A) UMAP displaying all cells of T cell lineage observed across the three KRC stages (early, intermediate, moribund) with a graph-based population delineation method applied to illustrate T lineage subpopulations. (B) T lineage UMAP with each cell color coded according to KRC stage (early = red, intermediate = green, moribund = blue). (C) Focused dot plot analysis displaying expression intensity and frequency of immune checkpoint genes in the regulatory T cell subpopulation. (D) Trial timeline schematic of KRC mice treated with either IgG isotype control antibody (n=6) or anti-CTLA4 antibody (n=6). Timing of antibody administration, MR imaging and planned animal sacrifice are all noted on the timeline. (E) Table displaying the results of the isotype control vs CTLA4 antibody treatment in KRC mice. Radiographic response: >30% decrease in tumor volume on MRI; radiographic stability: between +30% and -30% change in tumor volume; Radiographic progression: >30% increase in volume of tumor; Death: moribund or dead animal prior to planned sacrifice time point. (F) Axial T2 weighted MR images of longitudinally followed KRC mice treated with isotype control antibody (top row) and anti-CTLA4 antibody (bottom row).

### Cxcl5 is decreased in KRC cancer cells and is a putative mechanism for KRC indolence and its distinct TIME

To understand the basis for the unique TIME observed in KRC mice versus prototypal KPC tumors, we sought to determine if KRC cancer cells differentially expressed paracrine signals, relative to *Rnf43* intact mouse PDAC cell lines, which could play a role in mediating the observed immune landscape. To that end, conditioned media from two murine cell lines derived from moribund KRC tumors *versus* two murine cell lines derived from *Rnf43* intact PDAC cell lines were compared using multiplex mouse cytokine arrays (**Figure 7A**). In this 2 x 2 cell line analysis only one secreted factor was consistently different between the two groups; CXCL5 which was decreased in the KRC cell lines relative to the *Rnf43* intact cell lines. We then confirmed this finding by use of ELISA on the conditioned media, and added a third cell line for each group (**Figure 7B**). Quantitative PCR analysis was performed on these cell lines with concordant results indicating a potential transcriptional mechanism for low CXCL5 ligand levels in KRC cell lines (**Figure 7C**). Moreover, we compared *Cxcl5* expression in the cancer cell population of KPC and KRC scRNAseq datasets and confirmed low gene expression *in vivo* in the KRC GEMM compared to the KPC GEMM (**Figure 7D**). Notably, in our KRC scRNAseq dataset, the only cell population that expressed the CXCL5 receptor, CXCR2, was a population of MDSCs which first appeared in the KRC intermediate stage and expanded in the KRC moribund stage (**Figure 3A, B**). *Cxcr2* expression was noted in the MDSC population of both KRC and KPC GEMMs (**Supp Fig 9**). We also compared *CXCL5* gene expression level in a cohort of laser capture microdissected human IPMN samples concurrently profiled for *RNF43* mutation and expression, and found that expression was indeed lower in *RNF43* mutated patient samples (**Supp Figure 5B**). In order to understand the transcriptional mechanism whereby *Cxcl5* expression is decreased in KRC cells we performed ATAC-seq analysis which revealed a marked decrease in chromatin accessibility in the regions flanking the transcription start site of *Cxcl5* in KRC cell lines relative to the *Rnf43* intact PDAC cell lines (**Figure 7F**). Given the decreased chromatin accessibility at the Cxcl5 regulatory regions in KRC cell lines we asked if it would be possible to increase Cxcl5 expression in KRC cell lines by inhibiting histone deacetylates (HDAC). Indeed, treatment with the pan-HDAC inhibitor, panobinostat, resulted in a dose dependent increase in *Cxcl5* expression in KRC cell lines with 22-173-fold change in *Cxcl5* gene expression at the 50nM treatment dose versus the vehicle control (**Figure 7G**). Furthermore, to determine the role of decreased *Cxcl5* expression in the KRC GEMM we employed lentiviral transduction to stably over-expressed *Cxcl5* in KRC cell lines, resulting in protein levels comparable to that observed in the *Rnf43* intact PDAC cell lines. We then orthotopically implanted 10^5^ KRC-*Cxcl5* overexpressing cells or 10^5^ KRC-empty vector transduced cells in the pancreas of littermate mice resulting in an immunocompetent orthotopic mouse model. At the end of the study, we noted that the *Cxcl5* overexpressing KRC cell line resulted in pancreas weights >3-fold than those observed in the empty vector transduced KRC cell line (**Figure 7I**). Importantly, no notable tumor growth on the MRIs of the empty-vector transduced orthotopic injections were observed whereas clear tumors were observed in the *Cxcl5*-overexpressing KRC cell line orthotopic implants (**Figure 7J**).

**Figure 7:**
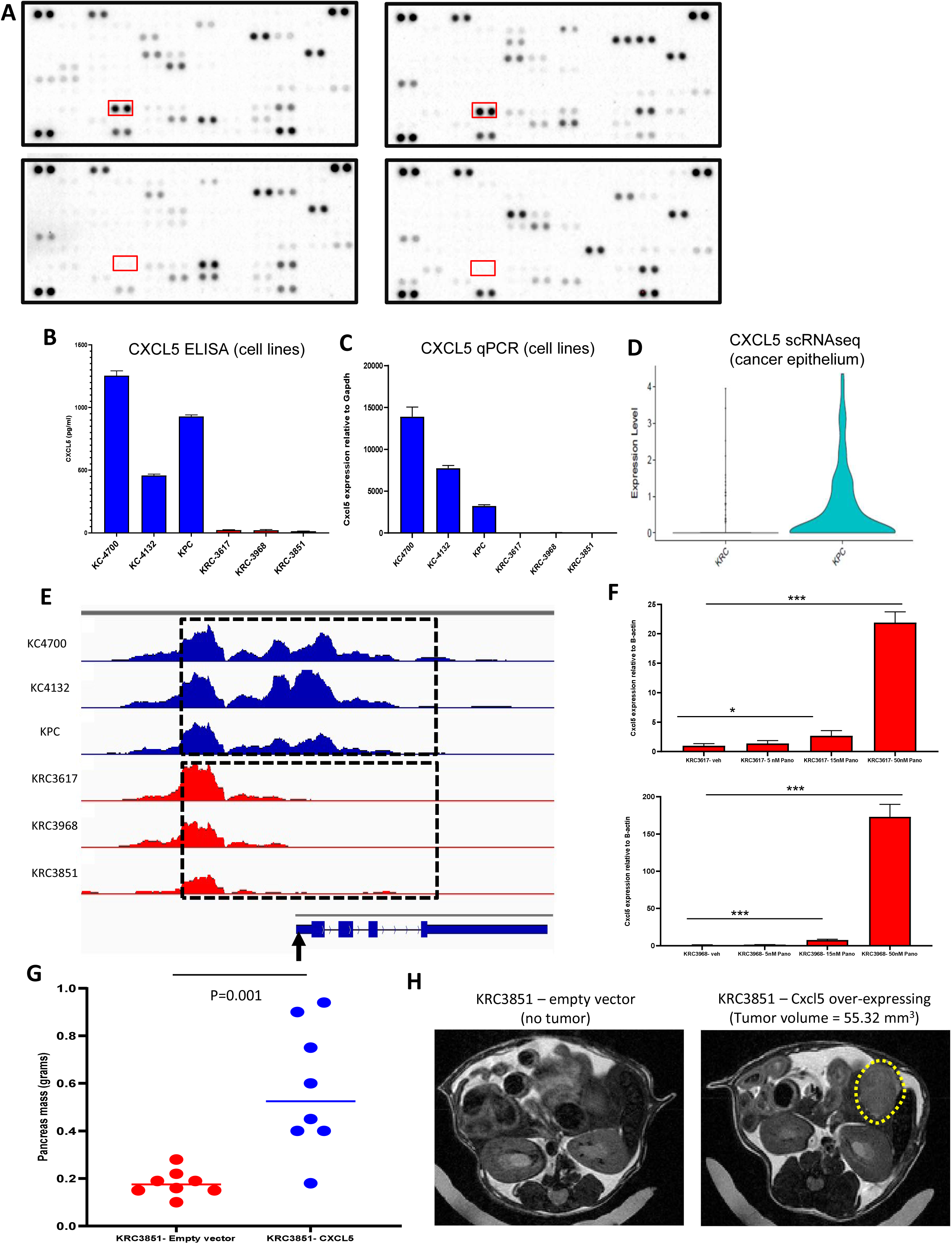
**CXCL5 is downregulated in the KRC cancer epithelium**. (A) Mouse cytokine arrays displaying relative intensities of proteins secreted into the media after 48 hour incubation with KC (top, left) and KPC (top, right) cell lines or KRC cell lines (bottom). The red rectangle outlines the dots corresponding to CXCL5. (B) ELISA confirmation of mouse CXCL5 on 48 hour conditioned media from 3 *Rnf43* wild type and 3 KRC mouse PDAC cell lines. (C) Quantitative PCR analysis of *Cxcl5* in the same 3 *Rnf43* wild type and 3 KRC mouse PDAC cell lines. (D) Violin plot of Cxcl5 expression scRNAseq datasets for KRC and KPC GEMMs. (E) ATAC-seq analysis of the transcriptions start site of *Cxcl5* in the 3 *Rnf43* wild type mouse PDAC cell lines and 3 KRC mouse PDAC cell lines. -3.47 log2 fold change (p= 5.52×10^-7^) between the *Rnf43* wild type and KRC mouse PDAC cell lines (regions outlined by the hashed boxes). Vertical arrow indicates transcription start site. (F) qPCR analysis of *Cxcl5* in 2 KRC mouse PDAC cell lines treated with increasing doses of Panobinostat. (G) Pancreas weights at 4 weeks in mice orthotopically injected with a KRC cell line transduced with empty vector (red) or *Cxcl5* (blue). 10^5^ cells/mouse were implanted in all experiments. The empty vector group had an average pancreas weight of 0.18g and the *Cxcl5* transduced group had an average tumor weight of 0.58g (p= 0.001). (H) Representative T2 weighted MRI (axial view) of mice orthotopically implanted with empty vector transduced KRC cells (left) and *Cxcl5* transduced KRC cells (right). (* p<0.05, **p<0.01, ***p<0.001)

## Discussion

In this study, we provide rigorous *in vivo* validation that *Rnf43* is a *bona fide* tumor suppressor gene in PDAC, and that it cooperates with mutant *Kras* in accelerating the development of precursor cystic lesions, as well as invasive adenocarcinomas, thus recapitulating the cognate multistep process observed in humans. Notably, the KRC^het^ mice, which featured loss of only one *Rnf43* allele, had a decreased survival equivalent to the KRC mice harboring loss of both *Rnf43* alleles. This observation suggests that haploinsufficiency, and therefore a decreased *Rnf43* gene dosage, may be sufficient to drive the phenotype we observed in this study. Indeed, in previously sequenced human PDAC samples, *RNF43* mutated tumors demonstrated an alteration at only one copy of *RNF43* while the other copy remained intact [6]. Further investigations of the KRC *versus* KRC^het^ cancer epithelium will be required to understand the mechanisms by which even monoallelic *Rnf43* loss promotes the transformation to neoplastic epithelium in the presence of oncogenic *Kras*.

The most conspicuous feature of the KRC GEMM reported in this study is the unique tumor immune microenvironment (TIME) we observed during KRC progression. At the KRC early time point we observed that myeloid cells were the most abundant cell type with lymphocytes comprising only a minority of cells. However, over the course of KRC progression from early to intermediate and moribund status, the myeloid:lymphocyte composition reversed, such that the majority of cells were of lymphocyte lineage by the moribund stage. Specifically, we noted a substantial increase in lymphocyte lineage cells at the KRC moribund stage where they comprised greater than three quarters of all cells in comparison to the KPC tumors, where lymphocytes cumulatively made up only 2.5%-18.5% of cells at this stage [16]. In addition, previous scRNAseq analyses of KPC have shown that myeloid cells comprise the majority of cells in moribund mice [16, 17]. In contrast, while most myeloid cells in the early and intermediate KRC stages were tissue resident and inflammatory macrophages, these cells were markedly diminished by the KRC moribund stage. Indeed, tissue resident macrophages have been shown to persist and expand within the pancreas over the lifespan of a mouse and depletion of this macrophage subset results in a marked decrease in tumor size in the KPC GEMM [21]. More specifically, it has been demonstrated in KPC mice that *Adgre1+* macrophages mediate T-cell exclusion from the PDAC microenvironment and that depleting this population of macrophages promoted T-cell tumor infiltration, resulting in tumor regression when combined with a CD40 agonist [32]. In addition, a separate study demonstrated that a small molecule inhibitor of CSF1R depleted inflammatory, tissue resident macrophages from the KPC tumor microenvironment and resulted in an improved T-cell response, decreased tumor size and improved animal survival [33]. These studies highlight the role of tissue resident macrophages in contributing towards a pro-tumorigenic, immune suppressive milieu within the PDAC TIME. Indeed, the paucity of tissue resident macrophages observed during KRC progression may contribute to the prolonged survival of the KRC GEMM in comparison to the KPC GEMM [34] and is a hypothesis that requires further exploration in subsequent studies of the KRC GEMM.

The KRC GEMM featured the expansion of early plasma cells (*Jchain^+^/Igm^+^*) along with evidence of affinity maturation toward mature plasma cells which had lower *Jchain* expression and specifically expressed several immunoglobulin genes. To our knowledge, this is a unique feature of the KRC GEMM with prototypal published models such as KPC and KIC not displaying the presence of an expanded B cell population and terminally differentiated plasma cell population [16, 17]. While we have shown that B-cell differentiation and terminal plasma cell maturation occur in the KRC GEMM, the functional implications of these cell populations remain to be explored. In a neoadjuvant immune checkpoint blockade (ICB) treatment trial of melanoma patients, B-cell gene expression signatures were found to be the most differentially expressed genes in the tumors of patients who responded to ICB versus those patients who responded poorly to ICB [35, 36]. Interestingly, these B-cells were found to be localized to intra-tumoral tertiary lymphoid structures. These findings also held true in analyses of another ICB sensitive cancer, renal cell carcinoma. Moreover, a separate study of melanoma patient samples, demonstrated that B-cell rich tertiary lymphoid structures predicted a good clinical outcome for patients treated with ICB [37]. In the KRC GEMM the *Ctla4* immune checkpoint gene was found to be over-expressed, in concert with several other immune checkpoint genes, with increasing intensity and frequency in the T_reg_ cell compartment. We have demonstrated the potential activity of anti-CTLA4 antibody treatment in the KRC GEMM. This study is the first to our knowledge which putatively demonstrates an intrinsic ICB sensitivity in a PDAC GEMM. These findings open a potential translational opportunity for ICB in *RNF43* mutated PDAC patients that will need to be validated in clinical trials.

We also identified a pivotal chemokine, CXCL5, which plays a role in mediating the cellular composition of the unique KRC TIME. *Cxcl5* expression was epigenetically regulated and could be increased by HDAC inhibition. These data suggest increasing cancer cell HDAC activity over the course of KRC tumor development which negatively regulates *Cxcl5* expression. Further studies aimed at dissecting dysregulated HDAC signaling pathways in the KRC GEMM are required. Importantly, CXCL5 is a chemokine which has been shown to be overexpressed in a subset of PDAC tumors and is associated with poor patient survival [38]. Given low *Cxcl5* expression in KRC, in contrast to *Rnf43* wild type PDAC cells, this supports the notion of KRC being relatively indolent tumor compared to KPC. CXCL5 overexpression in human PDAC has been shown to positively correlate with increased intra-tumoral, immunosuppressive macrophages and neutrophils [39]. We also identified a population of MDSCs which are present in both the KRC and KPC GEMMs and are the only population to expression the CXCL5 receptor, CXCR2. The role of CXCR2^+^ MDSCs in the KPC GEMM has previously been explored [40]; small molecule inhibition of CXCR2 resulted in increased intra-tumoral T-cell entry and sensitization to ICB which ultimately prolonged animal survival (**Supp Fig 9**). While these data demonstrate that CXCR2 inhibition is a strategy for ICB sensitization in the KPC GEMM, the low expression of the CXCR2 ligand, *Cxcl5*, in the KRC GEMM may render this model intrinsically sensitive to ICB

The lymphocyte-rich, macrophage-poor TIME seen in the KRC GEMM may, at least in part, begin to explain the observation that while the survival of KRC mice is shorter than KC mice, it is nonetheless improved in comparison to KPC (**Supp Table 1**) [19, 41]. The impact of *RNF43* mutations on the natural history of PDAC has not been studied in detail, although the higher frequency of alterations in precursor lesions (∼30%) compared to invasive adenocarcinomas suggests the possibility of a negative selection against early lesions bearing mutant *RNF43*; of note, in the 5-10% of PDAC patients in the TCGA dataset who harbor activating *KRAS* mutations and loss of function *RNF43* mutations, an improved survival is observed compared to *RNF43* wild type cases [42] **(Supp Figure 5C)**. Our data provides the underlying rationale to further expand on this observation and further leverage therapeutic dependencies such as susceptibility to immune checkpoint inhibitor therapy to further improve survival in this subset of patients.

## Supplementary Figures

**Supplementary Figure 1:**
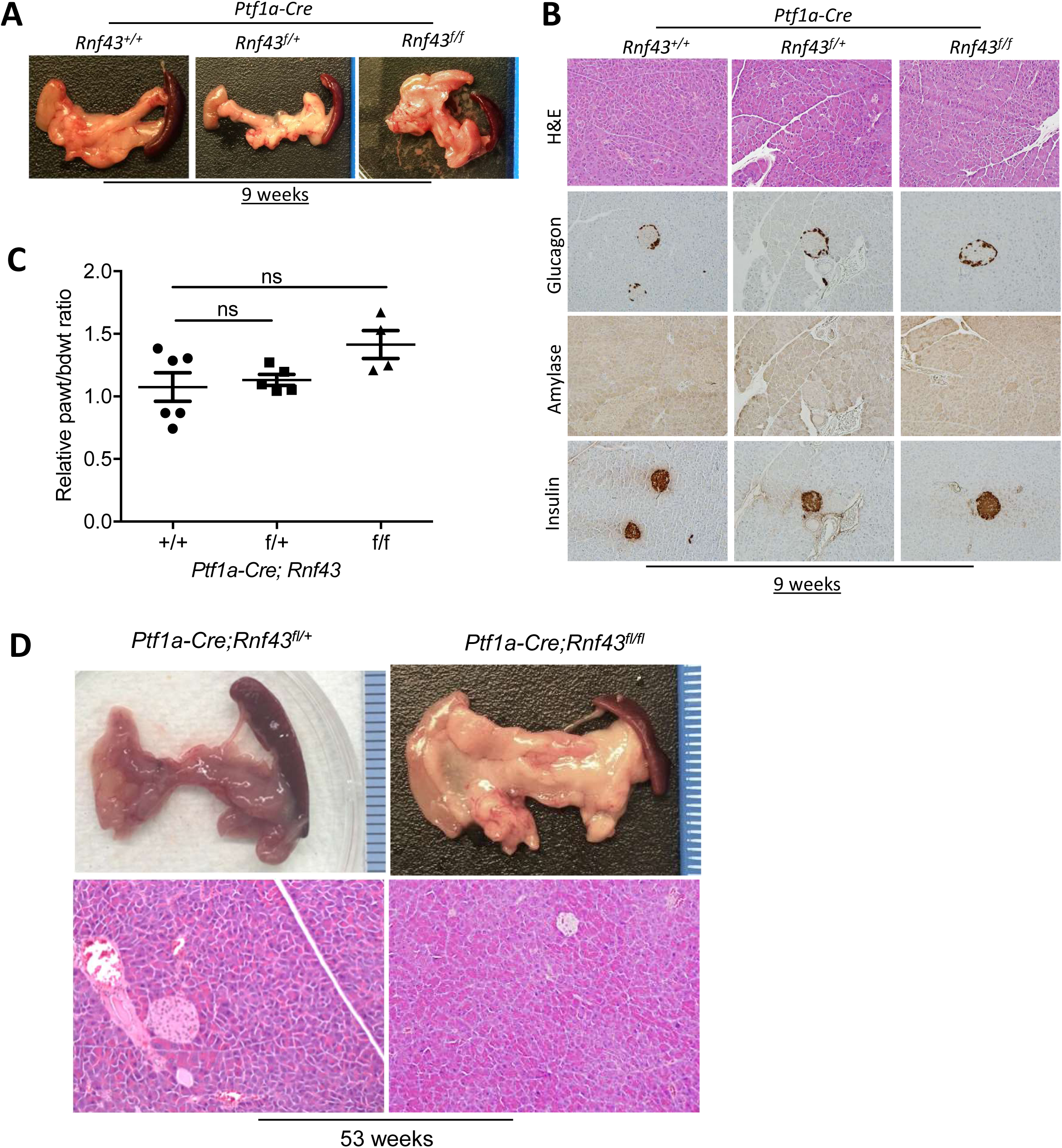
***Rnf43* is dispensable for normal pancreas development and insufficient for pancreatic neoplasia development.** (A) Representative images of gross morphology of pancreas of two-month old mice with conditional knockout of *Rnf43* alone without *Kras* mutation; (B) H&E staining of representative mouse pancreas at two months of age from indicated genotypes, 200X magnification (top), immunohistochemical staining of glucagon, amylase and insulin. (C) Relative pancreatic weight (paWt) to body weight (bdWt) ratio of control (*n* = 6), *Ptf1a*-*Cre*; *Rnf43^f/+^* (*n* = 5), and *Ptf1a-Cre*; Rnf43*^f/f^* (*n* = 5) pancreas at the age of two months. Student’s t-test was performed to calculate p-values; (D) Representative gross images of mouse pancreas at 53 weeks of age from indicated genotypes (top) and H&E staining of representative mouse pancreas tissue from indicted genotypes, 40X magnification (bottom).

**Supplementary Figure 2:**
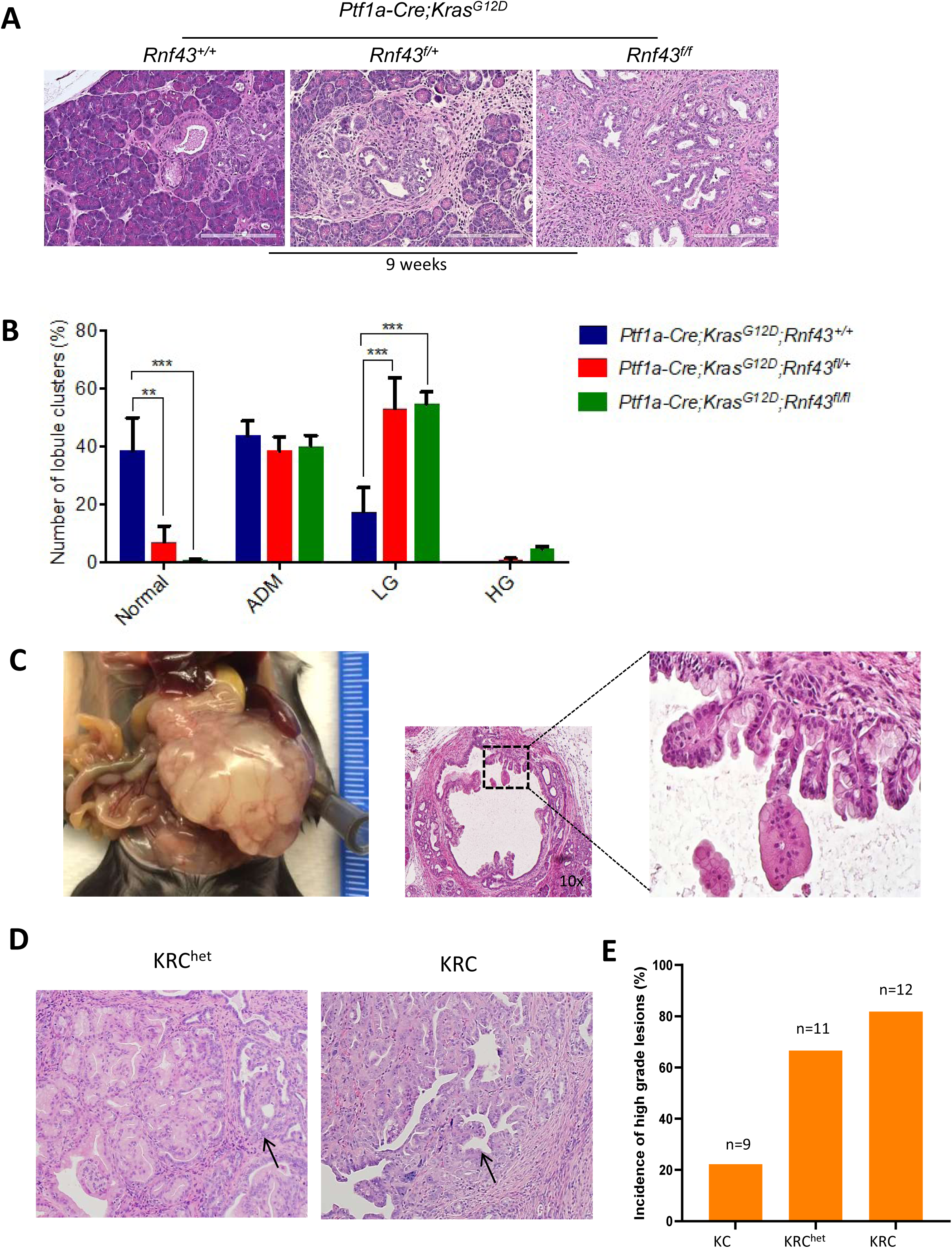
**Precursor lesions in Ptf1a-Cre;Kras^G12D^, *Rnf43* deficient mice progressed rapidly from low to high grade IPMN.** (A) Representative H&E tissue sections of pancreata from each of the three genotypes (KC, KRC^het^, KRC), showing lesions with increasing degree of dysplasia (40X magnification). (B) Graph representing the percentage of normal, ADM, low- and high-grade lesions in each genotype. 2-way ANOVA was used for multiple comparisons (**p<0.01; *** P<0.001, error bars denote the SEM). (C) Gross pancreas morphology (left) from a 3 month old KRC mouse demonstrating cystic components. Low magnification (center, 4X) and higher magnification (right, 10X) of the cystic lesion recapitulates human pancreatic IPMN. (D) H&E sections from mouse pancreata showing high-grade lesions (black arrow) (E) Graph showing percentages of mice below the age of 5 months with high-grade pancreatic precursor lesions.

**Supplementary Figure 3:**
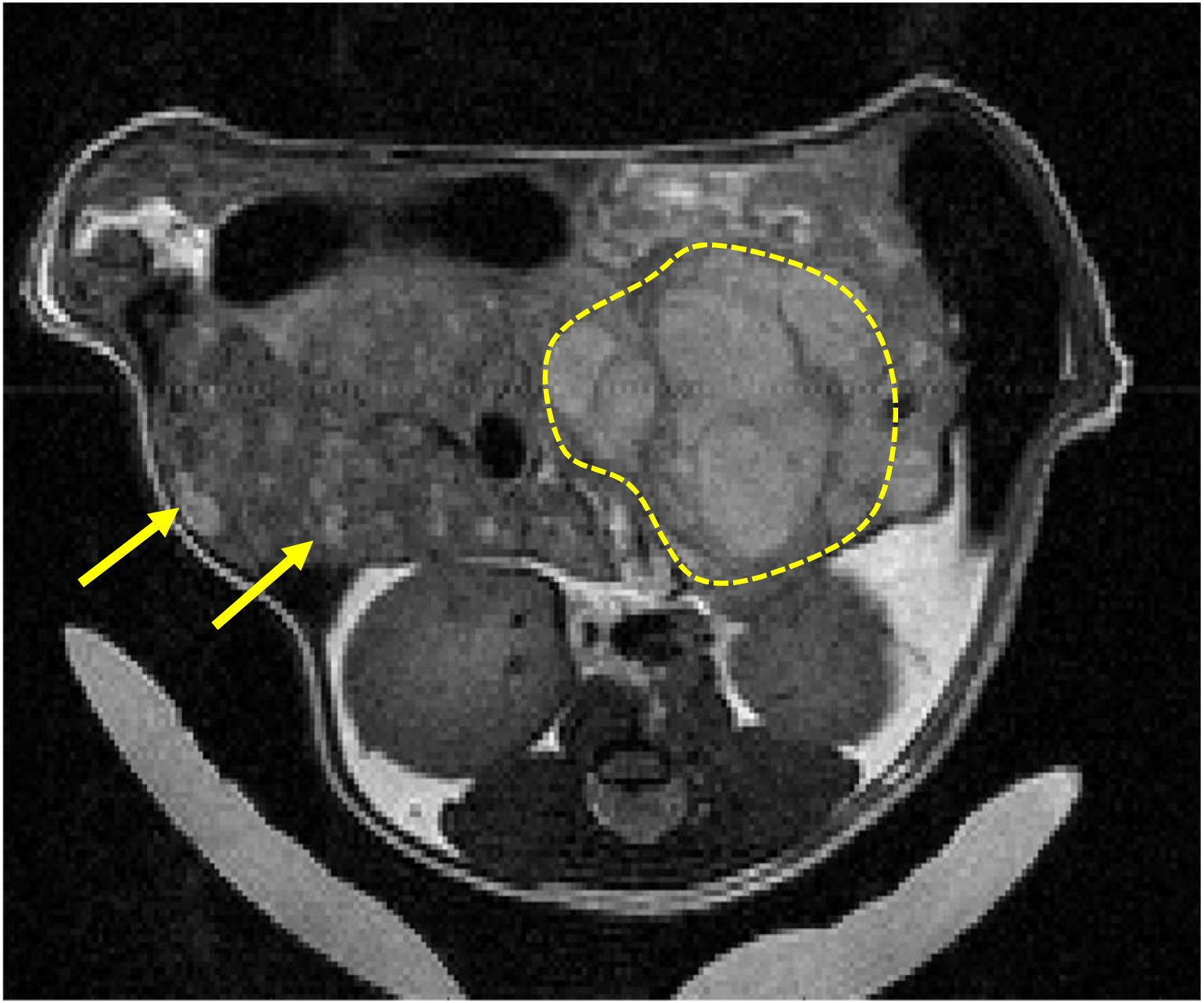
T2 weighted MRI of KRC mouse (axial view) demonstrating the presence of a large primary pancreatic tumor (hashed yellow outline) and the presence of multiple liver metastases (yellow arrows).

**Supplementary Figure 4:**
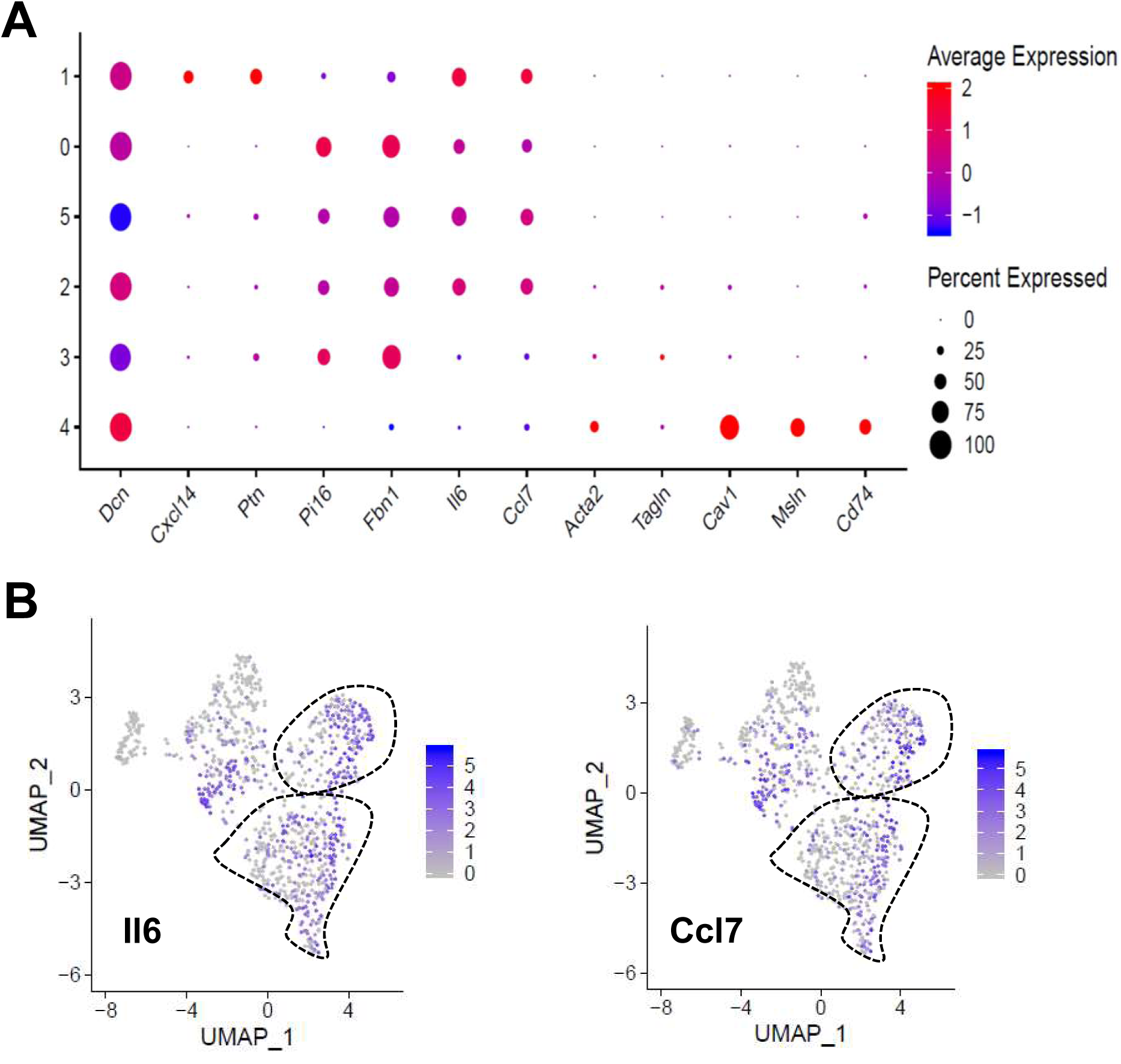
**Focused fibroblast analysis of the KRC GEMM.** (A) Dot plot of fibroblast markers applied to the sub-populations shown in the fibroblast UMAP. (B) Single gene UMAP for markers of inflammatory CAFs (Il6: left, Ccl7: right).

**Supplementary Figure 5:**
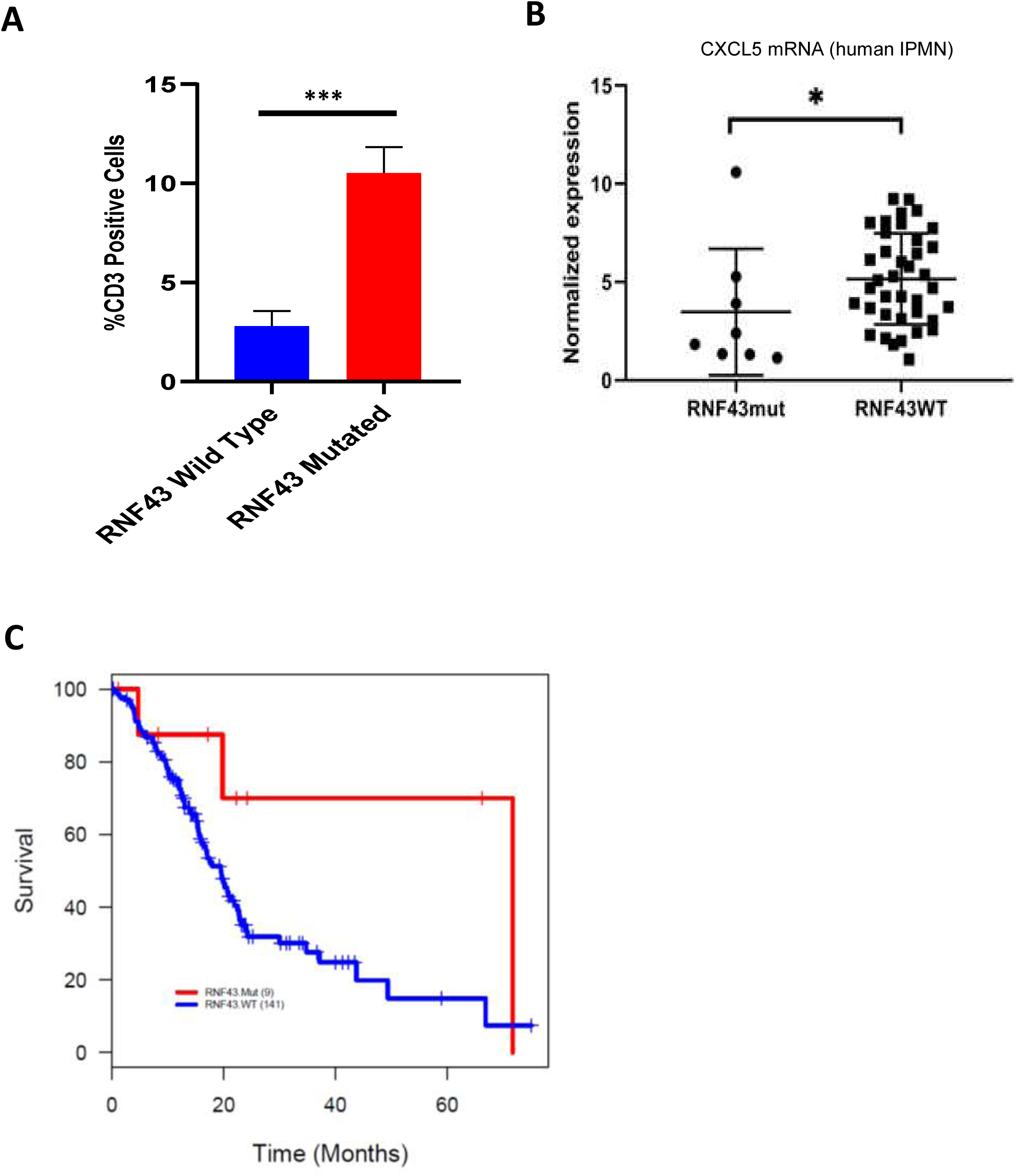
**Human *RNF43* mutated PDAC correlates.** (A) Comparison of T-cells (CD3^+^ cells) in *RNF43* wild type (n=6) versus *RNF43* mutant (n=4) PDAC patients, ***p-value=0.0005, paired t-test. (B) *Cxcl5* gene expression in a cohort of human IPMNs. *RNF43* mutated (n=8) and *RNF43* wild type (n=38) patients were compared. (*p-value=0.039, Mann-Whitney test) (C) Survival analysis comparing overall survival (OS) in the *RNF43* mutated versus the *RNF43* wild type PDAC patients from the TCGA dataset. A stratified Cox proportional hazards model for OS resulted in p-value=0.0479.

**Supplementary Figure 6:**
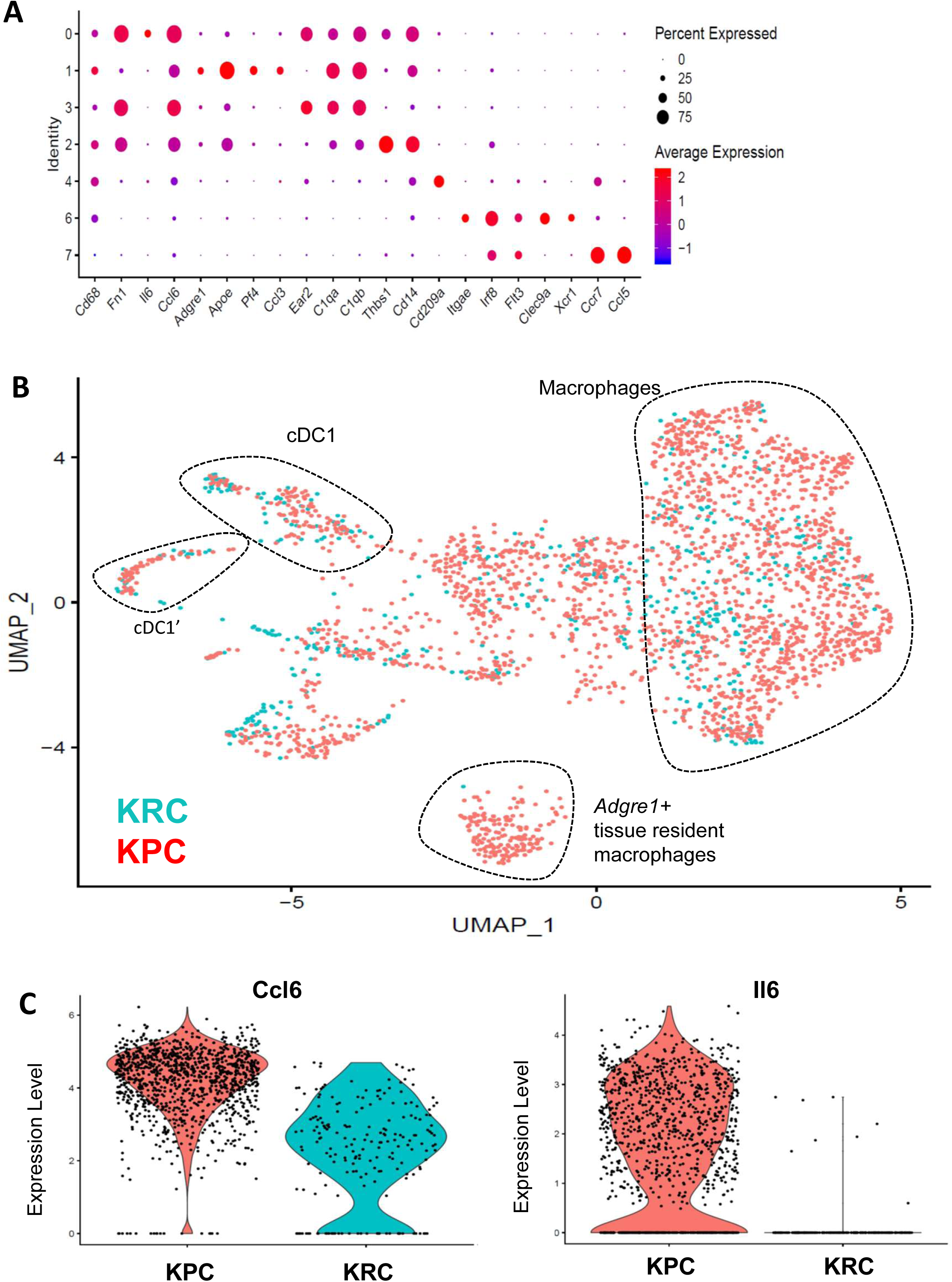
(A) Dot plot of myeloid markers applied to the sub-populations shown in the UMAP containing myeloid cells from early, intermediate and moribund KRC stages. (B) UMAP displaying all myeloid lineage populations in the KRC (teal) and KPC (red) GEMMs. *Adgre1^+^* tissue resident macrophages and the remaining macrophage populations are outlined. (C) Violin plots displaying cytokine expression (*Ccl6*: left, *Il6*: right) in KPC and KRC *Adgre1*-negative macrophages indicating a decreased relative inflammatory profile in the KRC macrophages GEMM.

**Supplementary Figure 7:**
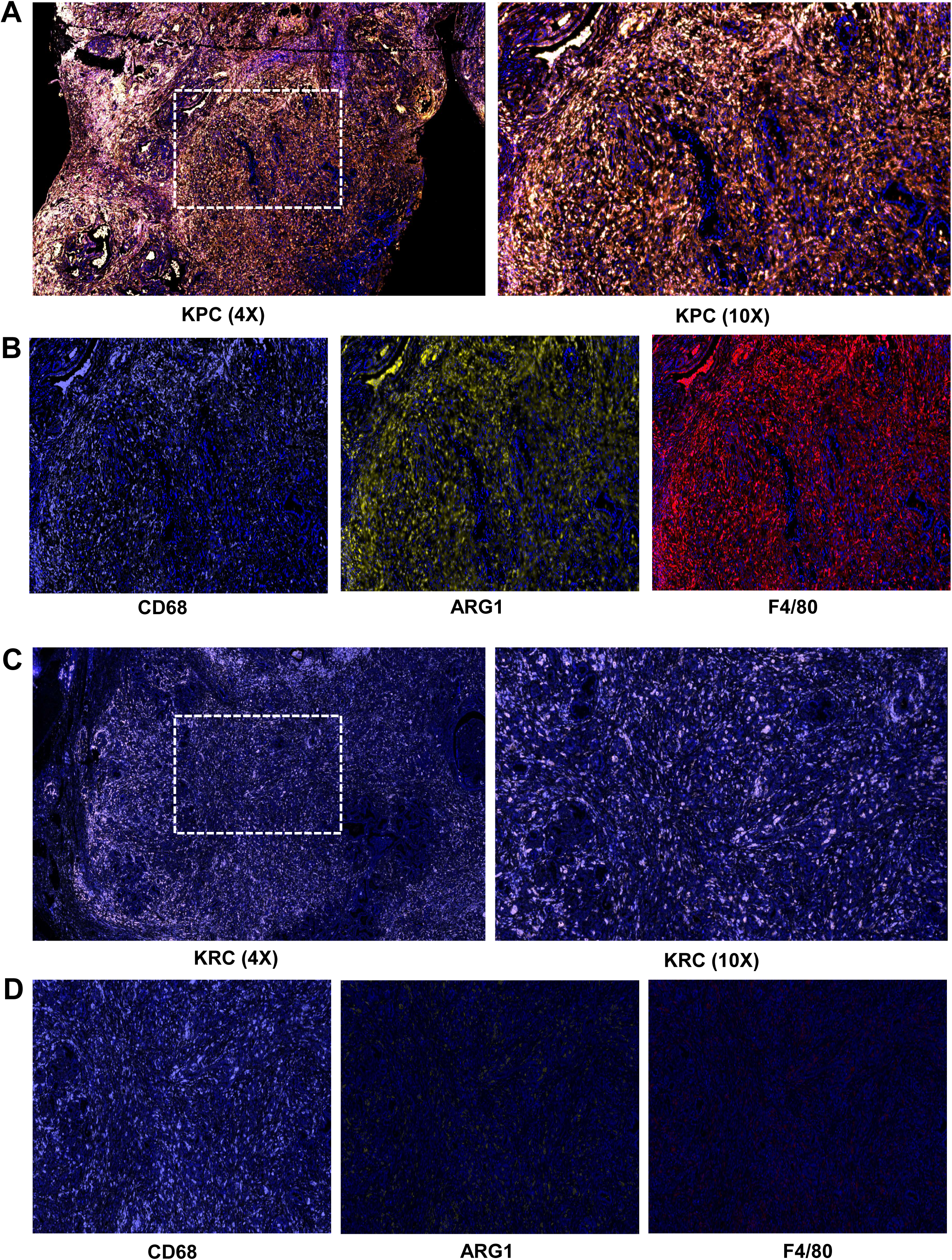
Immunofluorescence analysis (IF) of macrophage markers, CD68, ARG1, F4/80 in KPC and KRC tumors. (A) Three color IF of a KPC tumor with CD68 (white), ARG1 (gold), F4/80 (red) and DAPI (blue) displayed (4X magnification, left; 10X magnification, right). (B) Individual channels for CD68 (left), ARG1 (center) and F4/80 (right) displayed from the same KPC field at 10X. (C) Three color IF of a KRC tumor with CD68, ARG1, F4/80 and DAPI displayed (4X magnification, left; 10X magnification, right). (D) Individual channels for CD68 (left), ARG1 (center) and F4/80 (right) displayed from the same KRC field at 10X.

**Supplementary Figure 8:**
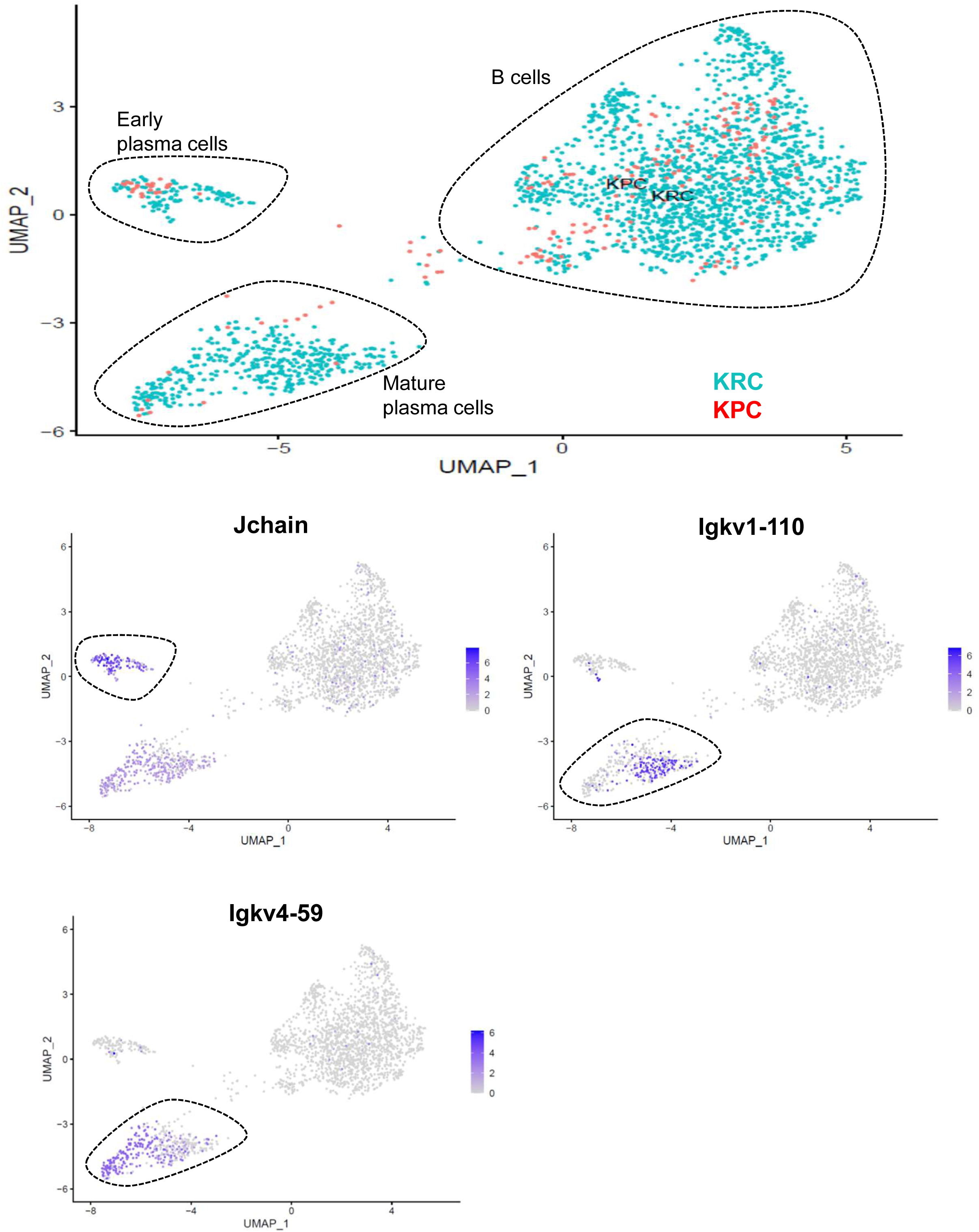
(A) UMAP displaying pooled B-cell lineage populations in the KRC and KPC GEMMs. Single gene UMAPs of the KRC-KPC B-cell lineage combined UMAP illustrating (B) *Jchain* expression, (C) *Igkv1-110* expression and (D) *Igkv4-59* expression.

**Supplementary Figure 9:**
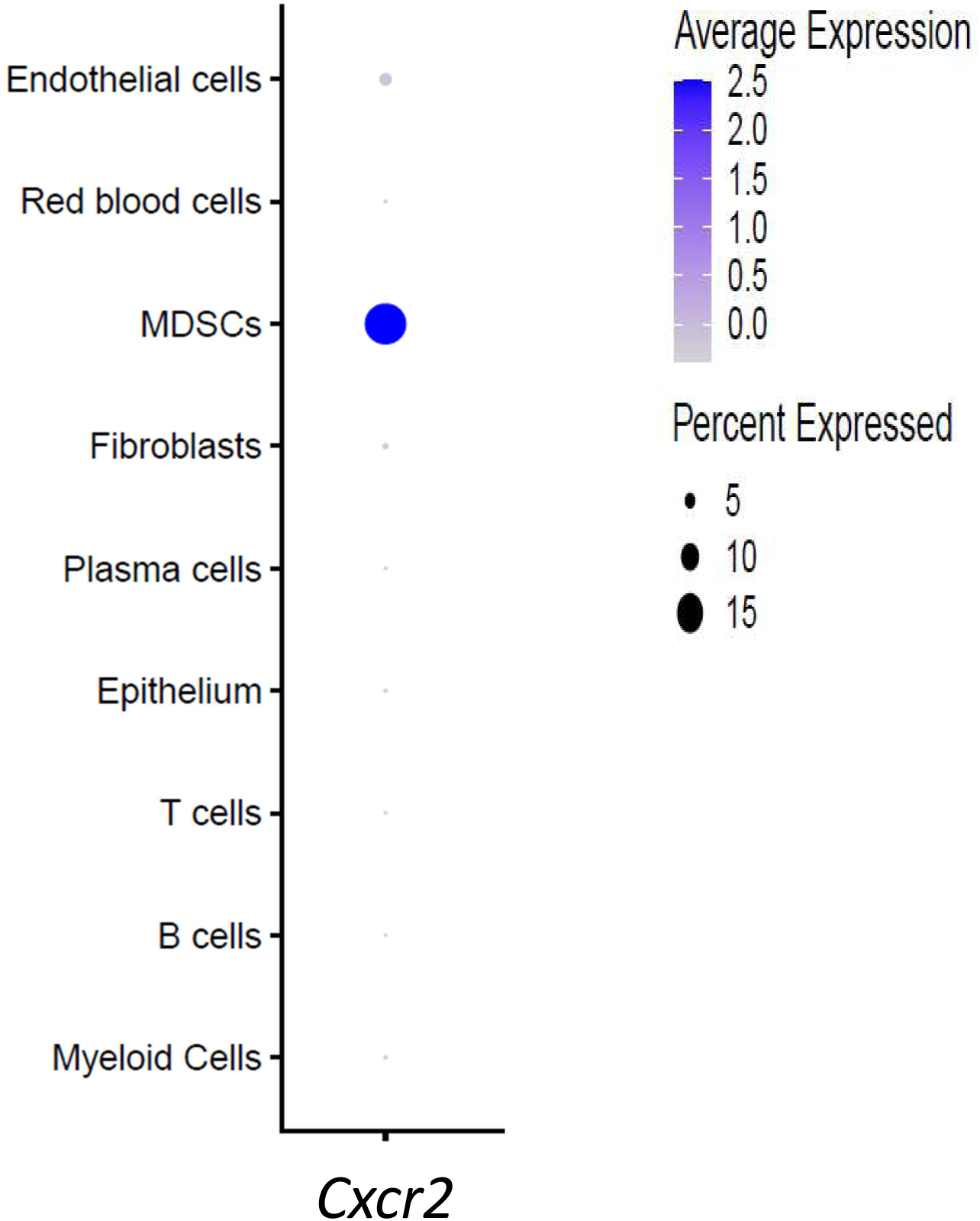
Dot plot displaying the expression of *Cxcr2* across major cell populations in the KRC-KPC pooled scRNAseq dataset. Legends denoting the Intensity and frequency of *Cxcr2* expression in each population is are displayed to the right of the dot plot.

**Supplementary Figure 10:**
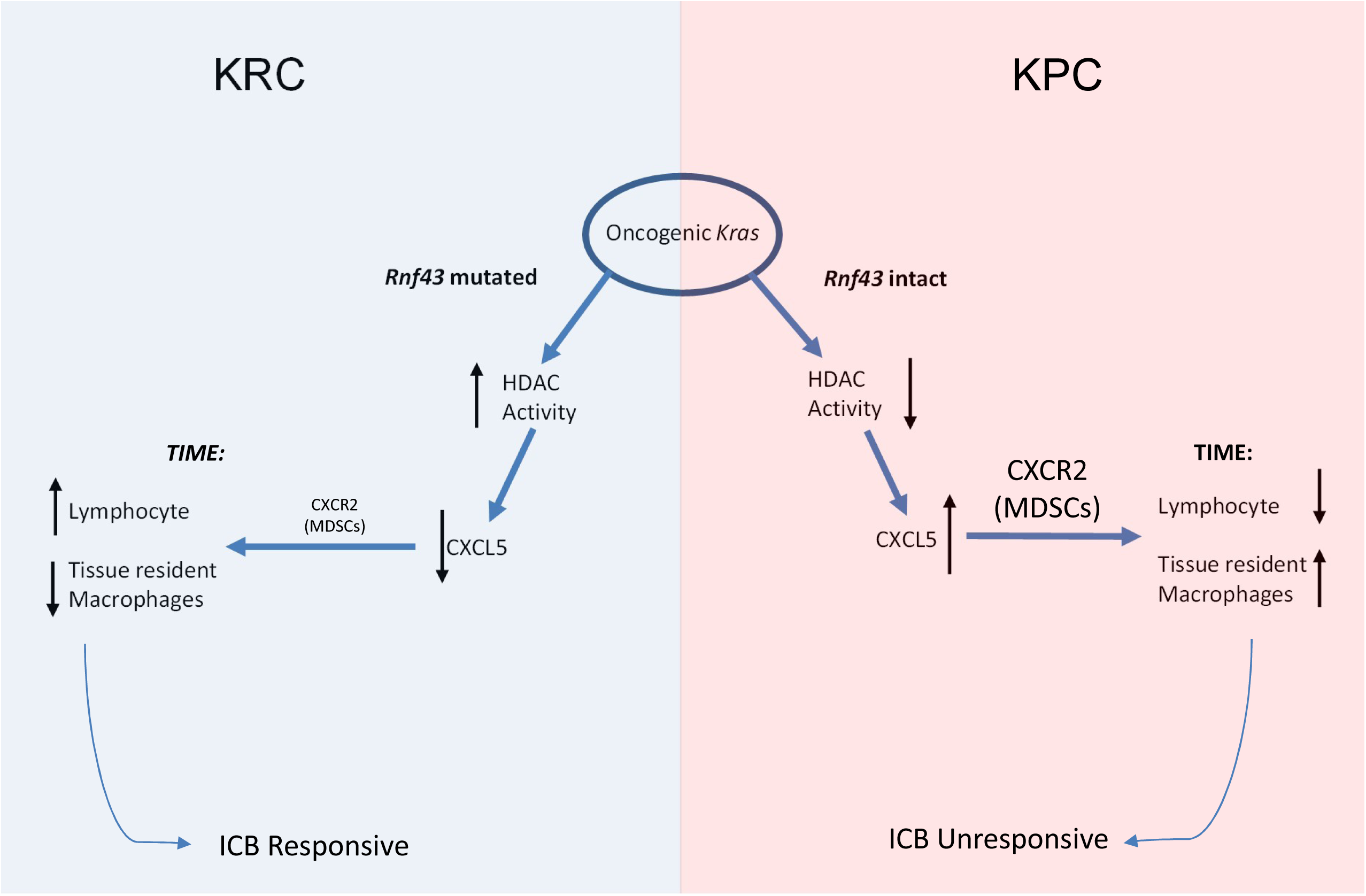
Schematic representation of the KRC tumor immune microenvironment (TIME) compared to KPC. Increased histone deacetylase (HDAC) activity in KRC leads to epigenetic downregulation of *Cxlc5* expression which culminates in an increased number of lymphocytes and decreased macrophages in the TIME. In turn, KRC tumors are sensitive to immune checkpoint blockade (ICB). PDAC GEMMs with oncogenic *Kras* and wild type *Rn43* do not display increased HDAC activity and *Cxcl5* expression is therefore not downregulated leading to increased stimulation of CXCR2^+^ MDSCs with the previously reported lymphocyte-poor, macrophage-rich TIME and lack of response to ICB.

**Supplementary Table 1:** Immune landscape and survival comparisons between moribund KRC and KPC GEMMs. Survival and each immune cell type as a percentage of total cells captured by scRNAseq analysis described in the study for the moribund KRC GEMM are displayed. The percentage of total cells for the KPC GEMM are listed as previously reported [16] as well as previously reported survival data [19].

|  | Moribund KRC | Moribund KPC |
| --- | --- | --- |
| Myeloid (%) | 12 | ~65 |
| T cells (%) | 21 | 2.5-18.5 |
| B cells (%) | 41 |  |
| Plasma Cells (%) | 15 | <1 |
| GEMM median survival (weeks) | 37 | 12-20 |

## Methods

### Generating conditional mice with pancreas-specific Rnf43 deletion

*Rnf43^fl/fl^* mice were obtained from the laboratory Dr. Hans Clevers (Hubrecht Institute, Netherland), and have been previously described ^27^. *Rnf43^fl/fl^* mice were crossed with the embryonic pancreatic epithelium restricted *Cre* recombinase strain, *Ptf1a-Cre*, or the derivative *Ptf1a-Cre;LSL-Kras^G12D^* mice (“KC”) to generate *Ptf1a-Cre;Rnf43^fl/+^, Ptf1a-Cre;Rnf43^fl/fl^, Ptf1a-Cre;Kras^G12D^;Rnf43^fl/+^* (“KRC^het^“) and *Ptf1a-Cre,Kras^G12D^;Rnf43^fl/fl^* (“KRC”) mice. Mice were maintained in C57BL/6 or DBA/2 genetic backgrounds. All mice were housed in pathogen–free barrier facility with food and water ad libitum. Animal studies were conducted in compliance with Institutional Animal Care and Use Committee (IACUC) guidelines of University of Texas MD Anderson Cancer Center. American Veterinary Medical Association Guidelines for the Euthanasia of Animals were strictly followed. Genotyping PCR was performed to confirm the genotype of mice using DNA from tails (Rnf43-FWD: 5’-GAAGCAGACAATGAAGCGAAT-3’, Rnf43-REV: 5’- TAGTGCCCCACAGAGGACA-3’, Kras1: 5’-GTCGACAAGCTCATGCGGGTG-3’, Kras2: 5’-AGCTAGCCACCATGGCTTGAGTAAGTCTGCA-3’, Kras3: 5’- CCTTTACAAGCGCACGCAGACTGTAGA-3’). For cell lines, recombination PCR was performed to confirm the *Cre*-mediated knockout of *Rnf43* in the pancreas (Rnf43-FWD: 5’- GAAGCAGACAATGAAGCGAAT-3’, Rnf43-REV: 5’ GAGGGTATCAGATCTCATTA- 3’, Kras1: 5’-GTCTTTCCCCAGCACAGTGC-3’, Kras2: 5’- CTCTTGCCTACGCCACCAGCTC-3’, Kras3: 5’- AGCTAGCCACCATGGCTTGAGTAAGTCTGCA-3’)

### Histology and immunohistochemistry

Mouse tissues from necropsies were fixed in 10% neutral buffered formalin and paraffin-embedded sections were stained with hematoxylin and eosin (H&E). Immunohistochemistry was performed as previously described [16]. Rabbit anti-Sox9 antibody (Millipore cat# AB5535) was performed at 1:500 dilution, rabbit anti-Ki67 antibody (CST cat# 12202) was performed at 1:400 dilution, rabbit anti-glucagon anitobody (CST cat# 2760) was performed at 1:100 dilution, rabbit anti-amylase antibody (Sigma cat# A8273) was performed at 1:400 dilution and rabbit anti-insulin antibody (CST cat# 3014) was performed at 1:400 dilution. For quantification of Ki67 positive cells, all immunostained nuclei independent of intensity were scored as positive. The areas that contain the largest number of Ki67 positive cells were selected. Positively stained cells were counted in 10 random fields per mouse at 1000X magnification. At least 3 independent mice per genotype were evaluated.

### Quantitative assessment of pancreatic lesions

To determine whether loss of *Rnf43* leads to acceleration of precursor lesions in the presence of a mutant *Kras^G12D^* allele, we performed necropsies of at least three mice per time point per genotype (“KC”, “KRC^het^”, and “KRC”) and compared the histology of age-matched mice from each cohort. Hematoxylin and eosin (H&E) stained whole tissue sections were scanned. The total lesional area per pancreas was manually delineated under bright field microscopic examination, and calculated using the Aperio ImageScope software (Leica Biosystems). Further, to quantify the prevalence of precursor lesion subtypes, a minimum of 45 acinar and/or ductal clusters was counted, and only the highest-grade lesion in each cluster was evaluated. Each cluster was assessed as normal, acinar-to-ductal metaplasia (ADM), low-grade (LG), or high-grade (HG) precursor lesion. This two-tier histological classification of pancreatic precursor lesions conforms to the revised classification system used in human. The data were expressed as a percentage of total clusters counted.

### Tissue digestion

Tumors and pancreas were enzymatically digested into a single-cell suspension as we have previously described [16]. Briefly, freshly dissected tumors were individually placed into a 10-cm tissue culture dish cut into fine pieces with a sterile blade. Samples were washed twice in PBS and added to 50 ml tube containing an enzymatic digestion buffer: collagenase type I (45 units/mL, Worthington Biochemical), collagenase type II (15 units/mL, Worthington Biochemical), collagenase type III (45 units/mL, Worthington Biochemical), collagenase type IV (45 units/mL, Gibco, Thermo Fisher Scientific), elastase (0.08 units/mL, Worthington Biochemical), hyaluronidase (30 units/mL, MilliporeSigma), DNAse type I (25 units/mL, MilliporeSigma) and 1% FBS. Tumors were incubated for 1 hour at 37 degrees Celsius. The cell suspension was washed three times with PBS and debris were filtered out using a 70-μm mesh filter (MilliporeSigma). Single cells were resuspended in 100 μL of PBS for subsequent downstream single-cell library preparation. Cell viability was measured by trypan blue. The viability of all samples was greater than 50%.

### Single-cell cDNA library preparation and sequencing

scRNAseq library generation was performed using the 10X Chromium System (10X Genomics Inc; Pleasanton, CA). Single cell suspensions were resuspended in PBS containing 0.04% *w/v* bovine serum albumin and brought to a concentration of 200–700 cells/μL. Appropriate volume of cells was loaded with single cell 5’ gel beads into a single cell chip and run on the Chromium Controller (10x Genomics, Inc; Pleasanton, CA). Dynabeads MyOne Silane magnetic beads (Thermo Fisher Scientific) were used to clean up the gel bead emulsion reaction mixture. Full-length, barcoded cDNA was amplified by PCR after cleanup. All samples were run on Agilent Tapestation 4200 using DNA high sensitivity D5000 tape. During library preparation, sample index and Illumina adapter sequences (Illumina Inc; San Diego, CA) were added. After library preparation quality control was performed using DNA D1000 tape on the Agilent Tapestation 4200 and final concentration was measured using the Qubit 4 Fluorometer DNA HS assay (Thermo Fisher Scientific). All samples were loaded at a concentration of 1.5 pM and run on the Illumina NextSeq500 High Output Flowcell. The run configuration was 26 base pairs (bp) × 98 bp × 8.

### Single cell RNA-seq analysis

scRNAseq data were analyzed as we had previously described [16]. Cell Ranger version 3.0.2 (10X Genomics) was used to process raw sequencing data. After sequencing and data processing scRNAseq datasets for each time point were computationally pooled for each time point. Individual cells from each time point were then displayed on a respective UMAP plot. Downstream analyses were carried out using Seurat (10X Genomics). KPC and KPfC data sets were obtained from our previously published report [16]. All scRNAseq data produced are to be deposited on the Gene Expression Omnibus platform (https://www.ncbi.nlm.nih.gov/geo)

### Immunofluorescence

Multispectral immunofluorescence analysis was performed on the Leica Bond RXm Processing Module fully automated staining platform (Leica Biosystems; Wetzlar, Germany). F4/80 rabbit polyclonal antibody (Abcam cat# 100790) staining was performed at 1:100 dilution followed by secondary antibody 520 reagent (1:100) (Akoya Biosciences; Menlo Park, California) and peak wavelength at 494/525. CD68 rabbit polyclonal antibody (Abcam cat# 125212) staining was performed at 1:100 dilution with secondary 620 reagent (1:100) (Akoya Biosciences) and peak wavelength at 588/616. Arg-1 rabbit polyclonal antibody (Invitrogen cat# PA5-85287) staining was performed at 1:15,000 dilution followed by secondary antibody 570 opal reagent (1:100 dilution) (Akoya Biosciences) and peak wavelength at 555/570. For human IF analysis, anti-CD3 Rabbit monoclonal antibody was used (Abcam, Ab16669). DAPI was the last reagent used for this assay with peak wavelength at 368/461 for nuclei staining.

### GEMM antibody treatment

Six month old KRC mice were enrolled onto the anti CTLA-4 treatment trial and treated with either anti-CTLA4 antibody (clone 9H10; Bio X Cell, Lebanon, NH) or *InVivoPlus* Syrian hamster IgG isotype control antibody (Cat# BP0087; Bio X Cell). All antibodies were diluted in *InVivoPure*™ pH 7.0 Dilution Buffer (Bio X Cell) and administered to mice. Mice were treated with 250 μg of antibody, twice per week by intraperitoneal injection.

### Cell culture of primary mouse PDAC cell lines

Tumors were dissected from GEMMs and minced into fine pieces using a sterile blade on a 10ml dish. The minced tissue was suspended in 20 ml of 1X Accutase Cell Detachment Solution (Thermo Fisher Scientific) and placed to rotate at 37 degrees Celsius for 4 hours. Cell suspensions were washed 3 times in RPMI containing 10% FBS with 1% penicillin/streptomycin (standard media). The cell suspension was then filtered through a 70-μm mesh filter and plated into a 24 well dish. Cells were allowed to adhere to the dish and expand for 1-2 weeks and then placed into a 10 cm dish for further expansion. Single cell colonies were obtained and cells were maintained in conditioned media in perpetuity.

### Magnetic resonance imaging analysis

MR imaging (MRI) was conducted in a 7T small animal MR scanner (Biospec®, Bruker Biospin MRI; Billerica, MA) using transmit/receive volume coils with 35 mm inner diameter. For MRI measurements mice were anesthetized using 2% isofluorane. For detecting tumors and assessing tumor volumes, morphologic MR images were acquired using a transversal T2-weighted turbo-spin echo sequence. The following parameters were used for all sequences: field-of-view of 4×3 cm^2^, matrix of 256×192 and spatial resolution of 156 micrometers, T2- weighted coronal rapid acquisition with relaxation enhancement (RARE) acquisition (T2-Cor, echo time, 38 ms; repetition time, 1800 ms; slice thickness, 0.75 mm; slice gap, 0.25mm) and a T2-weighted axial rapid acquisition with RARE acquisition (T2-Ax, echo time, 38 ms; repetition time, 3000 ms; slice thickness, 0.75 mm; slice gap, 0.25mm). For postprocessing, the tumor was segmented semi-automatically based on the T2-weighted MRI images, and the tumor volume was determined using an open-source image processing software (MATLAB, Mathworks; Natick, MA). The small animal imaging facility is supported by University of Texas MD Anderson Cancer Center Clinical and Translational Research Center which is funded by NCI #CA16672.

### Mouse cytokine array & ELISA

To assess cytokines secreted into the media of cell lines we allowed each cell line to reach 80-90% confluency over the course of 48 hours in 10ml of complete media in a 10cm cell culture dish. Supernatant was then collected and spun down at 180*xg* for 4 minutes, aliquoted into 1ml portions and stored at -80 degrees Celsius until further use. The Proteome Profiler Mouse XL Cytokine Array (Cat no: ARY028, R&D Systems; Minneapolis, MN) was used to assay 1ml of each supernatant per cell line assayed. The cytokine array procedure was performed precisely as instructed by the manufacturer. Cytokine arrays were developed and scanned on the ChemiDoc imaging system (Bio-Rad Laboratories; Hercules, CA). Individual dots were visually inspected for qualitative differences between groups and dots quantification confirmed by ImageJ software version 1.52k (National Institutes of Health, United States). Enzyme linked immunosorbent (ELISA) assay for mouse CXCL5 was performed on the supernatant collected for the cytokine arrays assay. The Mouse LIX Quantikine ELISA Kit (cat no: MX000; R&D Systems) was used per the manufacturer’s instructions.

### Quantitative PCR

RNA was using the by the RNeasy Kit (Qiagen) and reverse transcribed using the Superscript III cDNA Synthesis Kit (ThermoFisher Scientific). Quantitative PCR was performed using the SYBR Green Master Mix (ThermoFisher Scientific). Reactions were performed on a 7500 Real Time PCR System (Applied Bio Systems). Primers were designed in Primer3Plus web-based software. The following primers were used: Cxcl5_Fwd:GAAAGCTAAGCGGAATGCAC, Cxcl5_Rev: GGGACAATGGTTTCCCTTTT and Bet-Actin_Fwd: CCCTACAGTGCTGTGGGTTT, Beta-Actin_Rev: GACATGCAAGGAGTGCAAGA.

### Lentiviral transduction

*Cxcl5* (NM_009141) Mouse Tagged ORF Clone Lentiviral Particles containing 10^7^ transduction units/ml were purchased from Origene (Cat no: MR200761L4V; Rockville, MD). 50 μl of lentiviral suspension was added to sub-confluent KRC line in a single well of a 24-well plate containing 200 μl of complete media. An empty vector transduction experiment was performed in parallel. Cells were incubated in the lentiviral containing media for 72 hours and then media was aspirated and cells were transferred to a 6-well dish for expansion. Transduced cells were selected in puromycin containing complete media for 2-3 weeks at which time single cell clones were made. The supernatant of transduced cell lines were collected assayed by ELISA for mouse CXCL5 to confirm ligand levels comparable to KC and KPC cell lines.

### Orthotopic implantation

Littermates of the mice used for KRC cell line generation were used for orthotopic implantation. At approximately 4 months of age, mice were anesthetized with inhaled isoflurane followed by externalization of the pancreas. 10^5^ KRC cell lines (CXCL5 over expressing or empty vector containing) were resuspended in a 50μl Matrigel^TM^ solution (Corning Inc; Corning, NY) and injected into the body of the pancreas. The pancreas was returned to the abdomen and the fascia and abdominal wall was sutured. Buprenorphine was used analgesia after the procedure. Mice were monitored daily for signs of distress of excessive weight loss. All mice were sacrificed at 4 weeks and pancreata weighed, formalin fixed and paraffin embedded.

### ATAC-seq Analysis

ATAC-seq was performed following the Omni-ATAC protocol as previously described [43]. Briefly, live cells in culture were washed with phosphate buffered saline followed by harvesting and counting. Nuclei were isolated from approximately 50,000 cells in lysis buffer containing 0.1% NP-40 and 0.01% Digitonin and were fragmented with Nextera Tn5 Transposase (TDE1, Illumina) in TD Tagment DNA buffer (Illumina) for 30 minutes at 37 °C. The resulting library fragments were purified using a Qiagen MinElute kit. Libraries were further amplified 4–6 PCR cycles as described [43] and purified using AMPure XP beads (Beckman Coulter). ATAC-seq libraries were subsequently sequenced 2 x 75 bp on an Illumina NextSeq500.

## Notes

**Disclosures** A.M. receives royalties for a pancreatic cancer biomarker test from Cosmos Wisdom Biotechnology, and this financial relationship is managed and monitored by the UTMDACC Conflict of Interest Committee. A.M. is also listed as an inventor on a patent that has been licensed by Johns Hopkins University to ThriveEarlier Detection. All other authors are free of conflicts of interest and have nothing to disclose.

**Funding Sources** AM is supported by NIH grants R01 CA220236 and R01 CA218004. AM is also funded by the Sheikh Khalifa Bin Zayed Foundation

### Competing Interest Statement

The authors have declared no competing interest.

